# Ongoing brain state dynamics shape spontaneous awake reactivation and its relationship with memory

**DOI:** 10.64898/2026.09.24.754200

**Authors:** Hyun-Woong Kim, Ting Xu, Karl-Heinz Nenning, Alexandre Franco, Mark D’Esposito, Stan Colcombe, Michael Milham, Arielle Tambini

## Abstract

Memory reactivation is thought to support memory consolidation, but how ongoing brain states shape awake reactivation and its relationship to memory remain poorly understood. Here, we asked whether spontaneous hippocampal-cortical reactivation during awake rest is embedded in ongoing brain state dynamics by examining its coupling with whole-brain activity patterns and infra-slow global signal (GS) fluctuations, and whether these dynamics shape its relationship to memory retention. Reactivation preferentially occurred during an internally oriented brain state characterized by default mode network activation and dorsal attention network suppression, as well as states that reinstated encoding-related visual activity. Reactivation also clustered around the peak of infra-slow GS fluctuations and in high GS amplitude time windows, features previously associated with lower arousal. Importantly, the relationship between reactivation and memory depended on the ongoing brain state: distinct whole-brain activity patterns expressed during reactivation were associated with opposing memory outcomes, while reactivation during high GS amplitude windows was positively associated with retention. Together, these findings show that awake memory reactivation is embedded in dynamic whole-brain states that may influence its memory outcome, suggesting that recurring brain states may provide favorable windows for awake memory consolidation.

## Introduction

Spontaneous brain activity exhibits rich spatiotemporal dynamics across multiple timescales, ranging from infra-slow fluctuations spanning tens of seconds to brief sub-second events. One prominent example of low-frequency fluctuations is the recurrence of large-scale, whole-brain spatial patterns or ‘brain states’ which wax and wane on a timescale of seconds to tens of seconds (Bolt et al., 2022; Fox et al., 2005; Fransson, 2005; Gutierrez-Barragan et al., 2024; Liu et al., 2013; Smith et al., 2012). The cyclical emergence of such brain states, for example, those associated with internally versus externally directed processing (Fox et al., 2005), is in turn coupled with infra-slow fluctuations in arousal and global brain-wide activity (Raut et al., 2021; Yang et al., 2026; Bolt et al., 2025). Yet, this slow, spontaneous activity is punctuated by brief, transient events, such as the ‘replay’ or reactivation of neural activity patterns corresponding to recent and/or future experiences (Lee & Wilson, 2002; Foster & Wilson, 2006; Buzsaki, 2015; Zhang et al., 2025). While extensive literature has characterized how brief reactivation events unfold relative to ongoing brain dynamics during *sleep* (Diekelmann & Born, 2010; Klinzing et al., 2019; Rasch & Born, 2013 for reviews), emerging work has begun to probe how *awake* brain states are linked to reactivation or memory (Higgins et al., 2021; Huang et al., 2024; Yang et al., 2024, 2026; Wamsley et al., 2023). However, whether the brain states accompanying awake spontaneous reactivation help explain its link to memory remains unclear. Understanding this relationship may clarify why associations between awake reactivation and memory vary across contexts (Hoffman et al., 2024; Tal et al., 2024; Tanrıverdi et al., 2023) and inform interventions targeting awake reactivation, analogous to successful approaches during sleep (Ngo & Staresina, 2022; Carbone & Diekelmann, 2024). To gain insight into this question, here, we assess how spontaneous awake memory reactivation events are coupled with ongoing slow fluctuations in whole-brain fMRI activity, and whether this dynamic coupling is related to memory.

Spontaneous reactivation, or ‘replay’, of neural representations of past experience is a key mechanism thought to facilitate synaptic plasticity and thus support memory consolidation and retention (Buzsáki, 2015; Carr et al., 2011; Findlay et al., 2020; Tambini & Davachi, 2019; Buzsáki, 1996; Diekelmann & Born, 2010; Frankland & Bontempi, 2005; McClelland et al., 1995). Neurophysiological recordings robustly show that sequences of neural activity associated with prior awake experiences or future behavior are spontaneously represented (or ‘replayed’) during brief sharp-wave ripple (SWR) events, which predominantly occur during ‘offline’ states like sleep (Ji & Wilson, 2007; Peyrache et al., 2009; Rubin et al., 2022; Skaggs & McNaughton, 1996; Zhang et al., 2018) and awake restful periods (Diba & Buzsáki, 2007; Foster & Wilson, 2006; Liu et al., 2019; Pfeiffer, 2020). Neuroimaging approaches, such as functional magnetic resonance imaging (fMRI), have also revealed evidence of spontaneous memory reactivation, whereby spatial patterns of spontaneous resting activity become more similar to patterns elicited during prior experience, such as learning or encoding tasks (Schuck & Niv, 2019; Tambini & Davachi, 2013; Yu et al., 2024). Despite the limited temporal resolution of fMRI, temporally brief or transient reactivation events can be reliably identified through dynamic, timepoint-by-timepoint analyses (Deuker et al., 2013; Gruber et al., 2016; Schapiro et al., 2018; Schlichting & Preston, 2014; Staresina et al., 2013; see also Logothetis et al., 2012). Importantly, reactivation is associated with later memory via correlational evidence (Deuker et al., 2013; Dupret et al., 2010; Schapiro et al., 2018; Tambini & Davachi, 2013; Staresina et al., 2013) and causal manipulations (Ego-Stengel & Wilson, 2010; Girardeau et al., 2009; Fernández-Ruiz et al., 2019). Beyond memory, reactivation has also been linked to adaptive behaviors such as spatial planning and navigation (Pfeiffer & Foster, 2013) and model-based decision-making (Mattar & Daw, 2018; Wimmer et al., 2023).

Given the functional role of reactivation, a key question is which brain states and behavioral conditions shape when reactivation events occur. Addressing this question may help identify when interventions can most effectively influence reactivation and memory. Characterizing reactivation in relation to ongoing brain dynamics during sleep (Klinzing et al., 2019; Oyanedel et al., 2020; Staresina et al., 2015) has enabled closed-loop memory modulation approaches (Antony et al., 2018; Ngo et al., 2013; Ngo & Staresina, 2022; Nicolas et al., 2025; Papalambros et al., 2017). In contrast, less empirical work has characterized when spontaneous reactivation occurs during wakefulness. Awake reactivation is commonly observed during quiescent periods, including pauses in ongoing behavior or awake rest (Diba & Buzsáki, 2007; Foster & Wilson, 2006; Tambini & Davachi, 2013, 2019; Liu et al., 2019). Theoretical models similarly propose that hippocampal function switches between an externally oriented state supporting memory encoding and an internally-oriented state that supports retrieval and consolidation (Buzsáki, 1989; Honey et al., 2017; Mednick et al., 2011). Low acetylcholine is thought to bias hippocampal physiology toward the internally-oriented state supporting consolidation (Buzsáki, 1989; Hasselmo, 1999; Hasselmo & McGaughy, 2004; Vandecasteele et al., 2014). Since cholinergic activity varies with environmental novelty (Giovannini et al., 2001), external attention (Arnold et al., 2002), and arousal state (Collins et al., 2023; Neyhart et al., 2024), lower arousal and less externally oriented states may favor reactivation and consolidation. Consistent with this framework, quiet rest after learning can benefit memory retention compared to externally-oriented tasks (Brokaw et al., 2016; Craig et al., 2015, 2016, 2018; Dewar et al., 2014; Wamsley, 2022; but see Humiston et al., 2019) and relatively offline or internally oriented awake states are associated with greater benefits of rest (Wamsley et al., 2023; Wamsley & Summer, 2020) or reactivation on memory (Tambini et al., 2017). Together, this evidence motivates the hypothesis that dynamic fluctuations in brain state during wakefulness shape both when spontaneous reactivation occurs and how it supports memory.

Resting-state neuroimaging can be used to harness the rich temporal dynamics present in spontaneous brain activity to test how slow fluctuations in large-scale network configurations and global brain activity shape awake reactivation. Resting brain activity slowly fluctuates between recurring, large-scale spatial patterns, including internally vs. externally-oriented configurations characterized by opposing activation of the Default Mode Network (DMN) and Dorsal Attention Network (DAN), as well as activation of opposing poles of the sensorimotor-association (S-A) axis (i.e., the principal gradient of cortical organization; Margulies et al., 2016; Bolt et al., 2022; Fox et al., 2005; Gutierrez-Barragan et al., 2019, 2024). Based on the theoretical framework described above, an internally oriented brain state characterized by concurrent increases in DMN activity and reductions in DAN activity may be particularly conducive to awake reactivation, consistent with the DMN’s established role in internally focused and self-referential processing (Andrews-Hanna et al., 2014; Raichle, 2015; Smallwood et al., 2021; Mevel et al., 2013) and episodic memory retrieval (Huijbers et al., 2011; Vincent et al., 2006). Consistent with this idea, a recent MEG study showed that transient reactivation bursts coincided with brief, sub-second activation of the DMN and a parietal alpha network (Higgins et al., 2021). These findings raise the possibility that slow fluctuations in brain state dynamics may shape awake reactivation and help explain its functional relevance.

A complementary dimension of large-scale dynamics is provided by slow fluctuations in the global fMRI signal (brain-wide BOLD signal average). The occurrence of distinct whole-brain spatial activity patterns is temporally coupled with the phase of the ongoing global signal (GS) (Gutierrez-Barragan et al., 2019, 2024). Moreover, the GS is linked to arousal dynamics (Raut et al., 2021; Bolt et al., 2025), with GS peaks associated with transient electrophysiological signatures of arousal drops (Liu et al., 2018). Slow fluctuations in the amplitude of the GS, e.g., measured across minute-long windows, are also associated with arousal, with higher GS amplitude reflecting lower arousal (Liu & Falahpour, 2020; Gonzalez-Castillo et al., 2022; Wong et al., 2013). Based on the theoretical framework described above (Hasselmo, 1999; Wamsley & Summer, 2020), states of lower arousal during wake may be more favorable to reactivation and consolidation, consistent with higher hippocampal SWR rates during periods of pupil diameter constriction (McGinley et al., 2015; Yang et al., 2025). However, it remains unclear whether and how awake memory reactivation is coupled to ongoing fluctuations in the global fMRI signal, and whether reactivation during specific GS features (e.g., those associated with lower arousal) may be most beneficial for memory.

In the present study, we asked whether awake memory reactivation is a non-uniform or state-dependent process by characterizing not only when it occurs during wakefulness, but whether the ongoing whole-brain state in which it occurs shapes its relationship to memory. To address this, we analyzed an fMRI dataset in which the reactivation of incidentally encoded object-face stimuli was described (Tambini & D’Esposito, 2020). The extended post-encoding rest scans in this dataset (27 min in total; Fig. 1A) provided extensive sampling to characterize brain state and reactivation dynamics, while the inclusion of transcranial magnetic stimulation (TMS) additionally allowed us to test whether perturbing post-encoding brain activity altered the coupling between reactivation and global brain dynamics. We identified post-encoding reactivation events that occurred jointly across the hippocampus and a cortical site involved in processing the encoded stimuli, i.e., Lateral Occipital Cortex (LOC) (Fig. 1B-D; Schapiro et al., 2018; Tanrıverdi et al., 2023; Yu et al., 2024). To characterize how reactivation events are linked with dynamic fluctuations in global brain states, we estimated distinct whole-brain fMRI activity patterns that recur over time (Gutierrez-Barragan et al., 2024; Liu et al., 2013). To investigate relationships between reactivation and GS fluctuations, we focused on two features of GS dynamics previously linked with arousal: the phase of the GS (Bolt et al., 2025; Liu et al., 2018) and slow fluctuations in GS amplitude across minute-long time windows (Gonzalez-Castillo et al., 2022; Wong et al., 2013). Overall, we addressed three questions. First, we asked whether spontaneous reactivation is distributed uniformly across awake brain states or occurs preferentially during an internally oriented network configuration (high DMN, low DAN activity) and during GS features previously related to lower arousal (peak GS phase and high GS amplitude windows). Second, and central to the behavioral relevance of reactivation, we asked whether the brain dynamics accompanying reactivation are related to memory retention. Third, to assess the flexibility of coupling between reactivation and global brain dynamics, we asked whether perturbing post-encoding brain activity using theta-burst TMS modulated this coupling. To summarize these complementary measures and questions, Table 1 lists each ongoing brain state feature we examined, describing the analytic approach and key results: coupling with reactivation, modulation by TMS, and relationships with memory retention.

**Figure 1.**
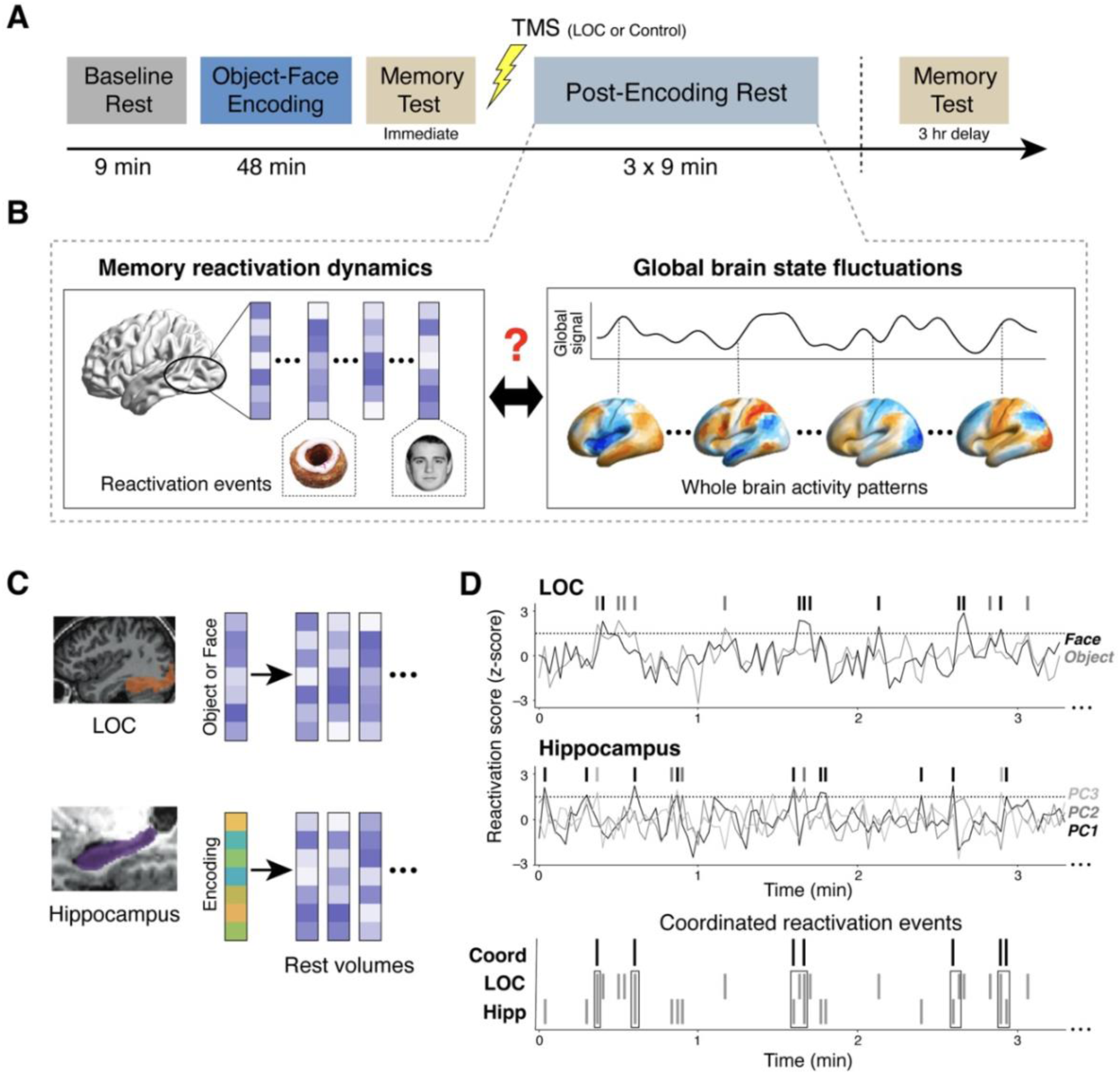
Experimental design and reactivation event identification. **(A)** Experimental procedure. Participants underwent baseline rest, object-face encoding, an immediate memory test, TMS (to LOC or a control site), post-encoding rest, and a delayed memory test. (B) Schematic of the central question. We asked whether memory reactivation dynamics during post-encoding rest (left; reactivation events reflecting reinstatement of encoded object or face patterns) are coupled with ongoing global brain state fluctuations (right; recurring whole-brain activity patterns and slow fluctuations in the global fMRI signal). (C) For LOC, template face and object patterns were derived from functional localizer scans. For the hippocampus, template patterns were derived from principal components (PCs) of encoding data. To identify reactivation events, we measured the similarity (correlation) of template patterns to the activity pattern at each post-encoding rest time point (TR), yielding the reactivation score time series shown in (D). (D) Example reactivation score time course for LOC and hippocampus (top, middle). Reactivation events (tick marks) were defined as time points exceeding 1.5 SD above the mean similarity (dotted line). Bottom panel: coordinated reactivation events (boxed) were identified as time points where hippocampal and LOC reactivation co-occurred at the same or an adjacent time point.

**Table 1.** Summary of analysis approaches and key findings.

| Feature | State | Method | Coupled with reactivation? | Coupling related to memory? | Coupling affected by TMS? |
| --- | --- | --- | --- | --- | --- |
| **Brain states**<br>(whole-brain activity patterns) | 1A (internally oriented) | k-means clustering (k = 8) of baseline-rest parcel time series | ✓ biased toward all four A poles (1A specific to reactivation vs. encoding) | ✗ | ✓ reduced coupling |
|  | 2A (visual) |  |  | ✓ greater coupling linked to *higher forgetting* | ✗ |
|  | 3A (unimodal sensorimotor) |  |  | ✗ | ✗ |
|  | 4A (visual + internal) |  |  | ✓ greater coupling linked to *higher retention* | ✓ reduced coupling; disrupted coupling–memory link |
| **GS phase** |  | Hilbert phase of band-pass filtered (0.01–0.05 Hz) GS | ✓ clustered near the GS peak | ✗ | ✗ |
| **GS amplitude** |  | SD of GS in 1-min sliding windows | ✓ more frequent in high-amplitude windows | ✓ linked to *higher retention* | ✗ |

## Results

### Memory Reactivation Event Identification

We identified spontaneous reactivation of recently encoded object-face stimuli during post-encoding rest as time points whose activity patterns showed high similarity to encoding-related template patterns (Fig. 1C-D), following prior work (Staresina et al., 2013; Schapiro et al., 2018; Tambini & D’Esposito, 2020; see Methods for more details). We focused on reactivation in two brain regions: the hippocampus, given its central role in associative memory formation and consolidation, and the LOC, as it represents object-face stimuli encoded in this study and shows robust post-encoding reactivation and memory-related hippocampal interactions in the same or nearby occipito-temporal regions (Tambini et al., 2010; Deuker et al., 2013; Schlichting & Preston, 2014; de Voogd et al., 2016; Murty et al., 2017; Kark & Kensinger, 2019).

We primarily focused on coordinated reactivation across the hippocampal-cortical network rather than events confined to a single brain region for two reasons. First, joint reactivation more directly reflects coordinated hippocampal–cortical activity, which is thought to support memory stabilization across distributed networks (McClelland et al., 1995; Maingret et al., 2016; Geva-Sagiv et al., 2023; Tompary & Davachi, 2017). Second, because the hippocampus and LOC tend to participate in distinct large-scale functional systems (Buckner et al., 2008; Grill-Spector et al., 2001), their conjoint reactivation is less likely to simply reflect activity within a particular large-scale network (e.g., hippocampal reactivation events co-occurring with DMN activation). Accordingly, we defined coordinated reactivation events as time points in which hippocampal reactivation was accompanied by LOC reactivation at the same or adjacent time point. This yielded an average of 42.1 coordinated reactivation events per participant (range: 27-63) across the three post-encoding rest scans.

To determine how these reactivation events were embedded within ongoing brain dynamics, we next examined three complementary features: recurring whole brain activity patterns, GS phase, and GS amplitude. For each measure of brain dynamics, we tested whether it was coupled with reactivation, whether this coupling was related to memory retention, and whether it was altered by TMS (Table 1).

### Brain State Identification

To examine whether awake memory reactivation occurs uniformly or preferentially during specific brain states (i.e., large-scale, whole-brain activity patterns), we characterized resting brain states and measured how frequently they occurred during reactivation events. We derived eight recurring spatial patterns of whole-brain activity from baseline (pre-encoding) rest scans, based on prior work showing that an eight-pattern solution reliably characterizes human resting state fMRI dynamics (Gutierrez-Barragan et al., 2024). To obtain a common set of whole-brain patterns across participants, we defined these spatial patterns at the group level. We used only baseline, rather than post-encoding, rest scans to avoid biasing brain state identification by reactivation of the specific memory representations being queried. As shown in Fig. 2A, the eight brain activity patterns formed four strongly anti-correlated pairs in both spatial topography and temporal expression. We thus conceptualized these as four brain states, with each state comprising two opposing poles of a single axis, labeled as ‘A’ and ‘B’ (e.g., Brain State 1 comprises the 1A and 1B poles). This organization aligns closely with prior work, as anti-correlated large-scale spatial patterns have been consistently observed across diverse analytical frameworks for characterizing brain dynamics (e.g., Bolt et al., 2022; Gutierrez-Barragan et al., 2024; Smith et al., 2012). Note that the assignment of ‘A’ and ‘B’ within each brain state was arbitrary; for ease of interpretation, we post hoc labeled the pole that occurred more often during reactivation events as the ‘A’ pole.

**Figure 2.**
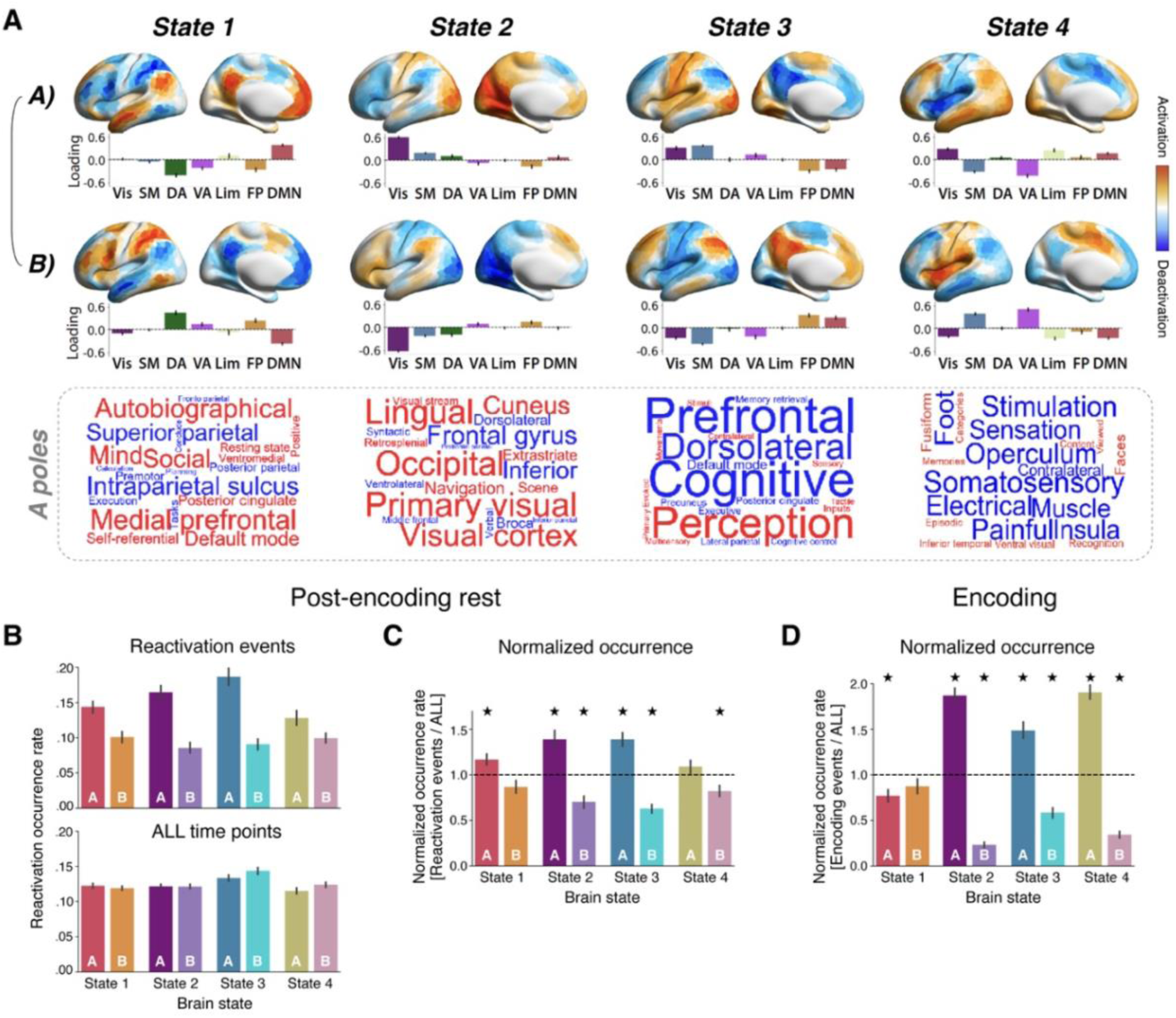
Brain states and their coupling with memory reactivation and encoding events. **(A)** Eight whole-brain activity patterns were derived from k-means clustering of baseline rest data. The eight patterns formed four strongly anti-correlated pairs in their spatial topography and temporal expression; each pair was conceptualized as a single brain state comprising two opposing poles, labeled ‘A’ (top row) and ‘B’ (bottom row). Cortical surface maps show the average spatial activation pattern for each pole. Bar plots below each whole-brain pattern show the mean activation within seven canonical large-scale networks (Yeo et al., 2011): visual (Vis), somatomotor (SM), dorsal attention (DA), ventral attention (VA), limbic (Lim), frontoparietal (FP), and default mode network (DMN). Word clouds show the top 10 positively (red) and 10 negatively (blue) associated Neurosynth meta-analytic terms for each ‘A’ pole, with text size representing the strength of association. (B) Occurrence rates for brain state pole (A and B) during post-encoding reactivation events (top) and across all post-encoding rest time points (bottom). (C) Normalized brain state occurrence rate during reactivation, obtained by dividing the occurrence rate of each pole during reactivation events by the occurrence rate across all post-encoding rest time points. Values above 1 (dashed line) indicate greater prevalence during reactivation, whereas values below 1 indicate reduced prevalence. (D) Normalized brain state occurrence during stimulus encoding, calculated analogously to reactivation but for encoding events. Stars in C-D indicate FDR-corrected one-sample t-tests against 1 (p < .05).

To qualitatively describe each brain state, we summarized the activity pattern of each brain state pole across seven large-scale cortical networks (Yeo et al., 2011; network loadings shown in Fig. 2A bar plots below brain maps). We also used a Neurosynth meta-analysis (Yarkoni et al., 2011) to visualize neuroscientific terms associated with each activation map (word clouds below brain maps in Fig. 2A). State 1 captured opposing patterns of putative internally oriented (1A) vs. externally oriented (1B) processing, characterized by differential weighting of the DMN, linked with internal processing, versus the dorsal attention network (DAN), linked with external processing (left bar plot in Fig. 2A) (Fox et al., 2005; Karahanoğlu & Van De Ville, 2015; Liu & Duyn, 2013; Vidaurre et al., 2017). This distinction was also reflected in the 1A word cloud, which exhibited positive terms related to internal processing (‘self-referential’ and ‘autobiographical’) and the DMN (‘medial prefrontal’ and ‘default mode’), along with negative terms related to external processing (‘tasks’ and ‘execution’) and DAN-related terms (‘intraparietal sulcus’). State 2 highlighted the visual network, which was positively weighted in 2A and negatively weighted in 2B (second bar plot in Fig. 2A). Similarly, the 2A word cloud prominently featured terms such as ‘visual cortex’, ‘occipital’, and ‘primary visual’. State 3 reflected the principal gradient of cortical organization, or sensorimotor-association (S-A) axis, differentially weighting unimodal primary sensory and motor cortices (visual and somatomotor networks) versus heteromodal association cortices (i.e., DMN and frontoparietal network, third bar plot in Fig. 2A) (Margulies et al., 2016; Nenning et al., 2023). Specifically, 3A captured the sensorimotor pole of the S-A axis, whereas 3B captured association pole. Lastly, State 4 contrasted visual, limbic, and default mode networks with somatomotor and ventral attention (salience) networks. Specifically, 4A positively weighted visual, limbic, and default mode networks while negatively weighting the somatomotor and ventral attention networks, whereas 4B showed the opposite pattern. Notably, the 4A word cloud included terms related to visual episodic memory processing (e.g., ‘fusiform’, ‘faces’, and ‘episodic’; Fig. 2A), suggesting that this state may be particularly relevant to object-face reactivation in this dataset.

### Brain State Classification during Reactivation

To identify the specific brain states linked with memory reactivation events, we classified individual time points during post-encoding rest as one of the eight isolated whole-brain activity patterns using a winner-take-all approach (i.e., labeling each time point as the specific brain state pole most similar to its activity pattern). Critically, because these whole-brain patterns were defined from the baseline (pre-encoding) rest data, any coupling between reactivation and particular patterns reflects a relationship between independently defined measures. As shown in the top plot of Figure 2B, reactivation events occurred non-uniformly during rest, with occurrence rates during reactivation significantly varying across the eight whole-brain activity patterns (repeated-measures ANOVA: *F*_7,357_ = 14.49, *p* < .001). Within each state, the pole occurring more frequently during reactivation was termed the ‘A’ pole, as described above.

To ensure that the differential occurrence of activity patterns during reactivation (Fig. 2B, top) was not driven by a non-uniform prevalence of specific brain activity patterns more generally, i.e., was not specific to reactivation per se, we measured the occurrence rate (frequency) of activity patterns across all time points (Fig. 2B, bottom). The occurrence rate was relatively uniform across all time points during post-encoding rest (Fig. 2B, bottom), with a small bias towards State 3 (reflecting the principal gradient). We directly tested whether occurrence rates differed between reactivation (Fig. 2B, top) versus all post-encoding rest time points (Fig. 2B, bottom) by calculating a normalized reactivation occurrence rate (the occurrence rate during reactivation events divided by all time points; Fig. 2C). Normalized occurrence rates that systematically differ from 1 indicate an increased or decreased prevalence of a particular brain state pole during reactivation per se. As in the prior analysis, normalized brain state occurrence rates during reactivation events varied significantly across brain state poles (*F*_7,357_ = 17.19, *p* < .001). Specifically, large-scale activity during reactivation was more likely to be labeled as 1A, 2A, and 3A, and less likely to be classified as 2B, 3B, and 4B (one-sample t tests: 1A: *t_51_* = 2.90, *p_FDR_* = .007; 2A: *t_51_* = 4.40, *p_FDR_*< .001; 3A: *t_51_* = 5.26, *p_FDR_* < .001; 4A: *t_51_* = 1.26, *p_FDR_* = .212; 1B: *t_51_* = –2.02, *p_FDR_* = .056; 2B: *t_51_* = –4.67, *p_FDR_* < .001; 3B: *t_51_* = –7.82, *p_FDR_* < .001; 4B: *t_51_* = –2.98, *p_FDR_* = .007; Fig. 2C). Increases in 1A occurrence during reactivation reflects DMN activation and simultaneous DAN deactivation (Fig. 2A, left column), suggesting reactivation occurred more frequently during an internally oriented processing state. Changes in State 2 prevalence (increases in 2A and decreases in 2B) show that reactivation was linked with visual network activation (Fig. 2A, second column). Changes in State 3 rates (increases in 3A and decreases in 3B) indicate that reactivation occurred more often during activation of the sensorimotor pole versus the association pole of the S-A axis (Fig. 2A, third column). Lastly, decreases in 4B prevalence suggest that reactivation may be reduced when visual, default mode, and limbic networks are deactivated along with activation of the ventral attention (or salience) and somatomotor networks (Fig. 2A, right column). These results demonstrate that reactivation does not occur uniformly during rest but is coupled with fluctuations in ongoing brain state dynamics.

We next asked whether the brain state configurations during reactivation were specific to ‘offline’ reactivation during rest or recapitulated those present during encoding. For example, we expect the prevalence of 1A, reflecting internally-oriented processing, to be specific to spontaneous reactivation in the absence of external stimulation (Higgins et al., 2021), whereas states reflecting stimulus-related visual processing, e.g., 2A, might be commonly expressed during encoding and resting reactivation. To compare the profile of brain states between reactivation and encoding, we isolated ‘encoding events’ (parallel to reactivation events during rest, i.e., time points strongly associated with stimulus encoding patterns during stimulus processing epochs) and classified each time point as a specific whole-brain activity pattern using the same procedures applied to rest scans. A two-way repeated measures ANOVA comparing normalized occurrence rates between encoding and reactivation events revealed a significant brain state pole × Encoding/Rest interaction effect (*F*_7,357_ = 22.3, *p* < .001), indicating that reactivation and encoding events showed distinct coupling with brain state poles (Fig. 2D). Specifically, 1A occurred significantly more often during reactivation than encoding (pairwise comparison: *t_51_* = 4.98, *p_FDR_*< .001). While 1A expression characterized resting reactivation (value > 1; Fig. 2C), it showed the opposite pattern (i.e., reduced prevalence) during encoding events compared to other time points during the encoding scan (value < 1; one-sample t test: *t_51_* = –3.52, *p_FDR_* = .001; Fig. 2D). This result suggests that an internally oriented state is selectively engaged during offline or spontaneous reactivation, while encoding is coupled with an opposing, externally oriented state.

In contrast to the crossover pattern observed for 1A, States 2-4 were modulated in the same direction during encoding and reactivation, consistent with reinstatement of brain states present during encoding. Specifically, during encoding, ‘A’ pole occurrence increased and B-pole occurrence decreased for all three states, largely mirroring the pattern present during resting reactivation (encoding: 2A: *t_51_* = 10.77, *p_FDR_* < .001; 3A: *t_51_* = 5.74, *p_FDR_* < .001; 4A: *t_51_* = 11.78, *p_FDR_* < .001; 2B: *t_51_* = –27.04, *p_FDR_* < .001; 3B: *t_51_* = –7.39, *p_FDR_* < .001, 4B: *t_51_* = –18.54, *p_FDR_* < .001). For States 2 and 4, brain state biases were significantly stronger during encoding than reactivation, with A poles occurring more frequently during encoding than reactivation (2A: *t_51_* = 4.24, *p* < .001; 4A: *t_51_*= 8.39, *p_FDR_* < .001) and B poles less frequently (2B: *t_51_*= – 6.65, *p* < .001; 4B: *t_51_* = –6.99, *p_FDR_* < .001), suggesting that reactivation recapitulates the directionality but not the full magnitude of encoding-related brain state expression. By contrast, State 3 occurrence did not significantly differ between reactivation and encoding events (3A: *t_51_*= 0.84, *p_FDR_* = .540; 3B: *t_51_* = –0.61, *p_FDR_* = .619). Together, these results indicate that resting reactivation is associated with two distinct patterns of brain state expression: an internally oriented processing configuration (1A) that is selective to spontaneous reactivation relative to encoding, and other brain states that are similarly expressed during encoding and reactivation, consistent with reinstatement of some whole-brain activity patterns during rest.

### Graded Brain State Strength during Reactivation and Functional Relevance to Memory

In addition to labeling each time point as a single brain state pole, we also characterized the graded brain state strength along each of the four brain state axes. This approach is advantageous because it provides continuous measures of brain state expression beyond categorical labels. To measure the graded brain state strength (Fig. 3A, top), we calculated the similarity (i.e., correlation) between the whole-brain activity at each time point and each of the eight brain state pole maps. Since the paired A and B poles capture opposing activity patterns, their graded expression or strength was, as expected, strongly negatively correlated over time (see Figure 3A for an example; all *R*s in each participant < –.96). We therefore merged strength values from the paired A and B poles into a single continuous measure for each brain state (A-strength minus B-strength divided by two; Fig. 3A) (Gutierrez-Barragan et al., 2024). Under this convention, positive brain state strength values indicate greater expression of the ‘A’ pole, negative values indicate greater expression of the ‘B’ pole, and zero indicates no bias toward either pole. To examine brain state strength during reactivation, we calculated the mean state strength across all reactivation events and subtracted the mean strength value across all post-encoding rest time points (analogous to normalized occurrence rates in Fig. 2C). For each of the four brain states, reactivation was associated with significant shifts toward the A pole (i.e., values > 0) (one-sample t tests: State 1: *t_51_* = 3.43, *p_FDR_* = .002; State 2: *t_51_*= 7.29, *p_FDR_* < .001; State 3: *t_51_* = 7.05, *p_FDR_* < .001; State 4: *t_51_* = 2.11, *p_FDR_* = .040; Fig. 3B), consistent with the prior classification results (Fig. 2B-C). Thus, the preferential coupling of reactivation with the A poles was also evident when brain state expression was measured continuously rather than categorically.

**Figure 3.**
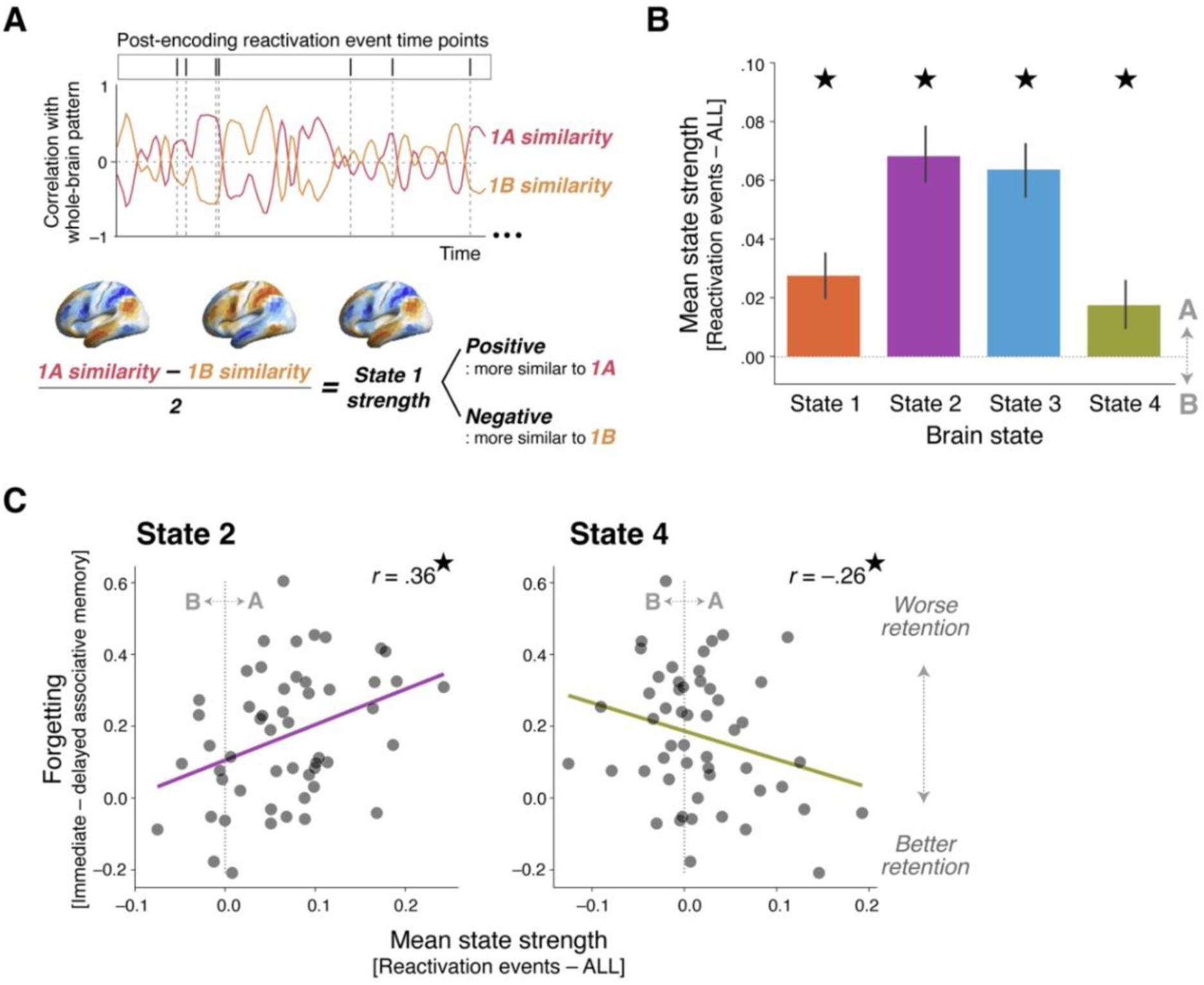
Graded brain state strength during reactivation and its relationship with memory. **(A)** Example time courses of similarity to the two opposing poles of a single brain state during post-encoding rest (1A, magenta; IB, orange). Similarity was calculated as the correlation between the whole-brain activity pattern at each time point and each pole’s spatial activity pattern. Reactivation event time points are marked by ticks and vertical dashed lines. For each brain state, a single graded strength value was computed at each time point as the ‘A’ similarity minus the ‘B’ similarity divided by two. Positive and negative values indicate greater strength toward the ‘A’ and ‘B’ poles, respectively. (B) Mean brain state strength during reactivation events, relative to mean strength across all time points, for each of the four brain states. Positive values indicate a shift toward the ‘A’ pole while negative values indicate a shift toward the ‘B’ pole. Stars indicate FDR-corrected one-sample t-tests against 0 (p < .05). (C) Scatter plots show the relationship between brain state strength during reactivation (relative to mean strength across all post-encoding time points) and associative forgetting (immediate minus delayed associative memory accuracy) for States 2 and 4. Each data point represents one participant.

To probe whether brain state expression during reactivation is stable or sensitive to perturbation, we examined whether theta-burst TMS to LOC (shown to reduce memory retention in this dataset, Tambini & D’Esposito, 2020) altered the coupling between reactivation events and ongoing brain states. A mixed-effects ANOVA on brain state strength during reactivation showed a significant interaction effect between brain state and TMS site (*F*_3,153_ = 4.12, *p* = .008; Fig. S1A). Pairwise comparisons indicated that TMS to LOC, relative to Control TMS, reduced brain state strength for State 1 (*t_50_* = 2.22, *p* = .031) and State 4 (*t_50_* = 2.22, *p* = .031) and marginally increased State 3 strength (*t_50_* = –1.95, *p* = .056) during reactivation events. That is, after LOC compared to Control TMS, reactivation events were no longer biased toward the A poles of States 1 and 4, and occurred during greater expression of the 3A brain state pole. These findings show that links between reactivation and ongoing brain states are not fixed, but instead are malleable and are sensitive to perturbation of post-encoding brain activity.

To test the behavioral relevance of brain state expression during reactivation, we investigated whether brain state strength during reactivation is related to associative memory retention, which is closely linked to hippocampal function and memory consolidation mechanisms (Davachi, 2006; Mayes et al., 2007; Tambini et al., 2010; Gruber et al., 2016; Murty et al., 2017). As a behavioral index of consolidation, we focused on changes in associative memory over time by calculating forgetting, inversely related to retention; higher values indicate greater forgetting or poorer retention. We first examined relationships between retention (forgetting) and brain state strength during reactivation across all participants, regardless of TMS site. We used multiple regression to probe each state’s unique relationship with memory while accounting for shared variance across brain states. The strength of States 2 and 4 during reactivation was significantly related to forgetting across all participants (Fig. 3C; State 2: *b* = 1.42 ± .52, *t* = 3.03, *p* = .004; State 4: *b* = –1.37 ± .52, *t* = –2.62, *p* = .012), whereas no reliable relationships were found for State 1 (*b* = –.16 ± .50, *t* = –0.32, *p* = .751) or State 3 (*b* = –.61 ± .52, *t* = –1.19, *p* = .241). Specifically, greater expression of the 4A brain state pole during reactivation was associated with less forgetting or better retention across participants (Fig. 3C, right). The opposite pattern was found for State 2: higher 2A state pole strength during reactivation was associated with more forgetting (Fig. 3C, left). These results suggest that expression of a brain state dominated by the visual network during reactivation was associated with poorer memory retention (as reflected by State 2), whereas a broader whole-brain pattern characterized by concurrent activation of visual, limbic, and default mode networks together with deactivation of ventral attention and somatomotor networks was associated with better retention (as reflected by State 4).

Since TMS influenced both memory retention and brain state expression during reactivation, we tested whether these relationships could be explained by, or differed across, TMS conditions. We repeated the analyses while modeling the TMS site as a between-group effect (mean-centered) and its interaction with brain state strength. For State 2, consistent with the results across all participants, brain state strength during reactivation remained significantly related to forgetting when TMS was modeled (main effect: *b* = 1.09 ± .34, *t* = 3.20, *p* = .002) and the TMS x State 2 interaction was not significant (*p* = .776), indicating that this relationship did not reliably differ between TMS sites. In contrast, the main effect of State 4 strength during reactivation was not significant when TMS site was modeled (*b* = –.29 ± .42, *t* = –0.70, *p* = .486), but the TMS × State 4 interaction was significant (*b* = 2.07 ± .84, *t* = 2.48, *p* = .017), indicating that the relationship between forgetting and State 4 strength during reactivation differed between TMS sites. State 3 showed similar, trend-level evidence for a TMS site-dependent relationship with forgetting (*p* = .052). For both States 4 and 3, relationships between brain state strength and memory were stronger in the Control TMS group (see Supplementary Results). Full interaction results and within-group correlations are shown for all four brain states in the Supplementary Results and Fig. S1B. Together, these results indicate that the relationships between brain state expression during reactivation and memory retention were not driven by pooling across TMS conditions, while also suggesting that the behavioral relevance of specific brain states, particularly State 4, is flexible and can be altered by perturbation of post-encoding brain activity.

### Functional Relevance of Overall Brain State Strength during Post-Encoding Rest

Previous work has shown that quiet rest after learning leads to better memory retention than engaging in externally oriented tasks (Brokaw et al., 2016; Craig et al., 2014; Dewar et al., 2014), and individuals who undergo brief ‘offline’ states more frequently exhibit better retention (Wamsley & Summer, 2020; Wamsley et al., 2023). These findings raise the question of whether a greater prevalence of internally oriented, as opposed to externally oriented, brain states during post-encoding rest might generally benefit memory retention, regardless of their coupling with reactivation events. To test this possibility, we asked whether the mean strength of each of the four brain states, averaged across the entire post-encoding rest period (i.e., rather than specifically during reactivation events), was related to associative memory retention. No brain state showed a significant association between its overall strength across all post-encoding time points and associative forgetting (State 1: *b* = –5.82 ± 8.06, *t* = –0.72, *p* = .474; State 2: *b* = –7.69 ± 6.23, *t* = –1.23, p = .224; State 3: *b* = 3.67 ± 8.15, *t* = 0.45, *p* = .654; State 4: *b* = 12.62 ± 7.05, *t* = 1.79, *p* = .080). Notably, for States 2 and 4, the direction of these associations, while not significant, was opposite to that observed during reactivation events. Together, these findings suggest that memory retention is not simply related to the overall prevalence of particular brain states during post-encoding rest, but instead to the brain states during reactivation.

### Global Signal Dynamics Associated with Reactivation

Complementing the above brain state analyses, we next examined two features of infra-slow GS dynamics that have been linked to ongoing arousal levels, GS phase and amplitude (Table 1; Gonzalez-Castillo et al., 2022; Liu et al., 2018; Wong et al., 2013). Specifically, we asked whether reactivation events are coupled to instantaneous GS phase, which varies over tens of seconds (Liu et al., 2018), and fluctuations in the GS amplitude across minute-long windows (Gonzalez-Castillo et al., 2022). To assess the stability and functional relevance of these relationships, we asked whether TMS modulated reactivation-GS coupling and whether coupling was related to associative memory retention.

To probe links between reactivation and GS phase, we extracted slow GS fluctuations (0.01-0.05 Hz), as in prior work (Raut et al., 2021), and compared them with reactivation events, as shown in an example fMRI scan in Figure 4A. We found that reactivation events clustered around the peak phase of the GS (i.e., 0°; Rayleigh test: *p* < .001; Fig. 4B). This phase locking did not differ by TMS site (Kuiper two-sample test: *V* = 0.036, *p* = .959; Fig. S2A), suggesting that GS phase coupling with reactivation reflects a stable rather than malleable feature of reactivation dynamics. Based on this finding, we tested whether the degree to which the GS was concentrated around the peak phase during reactivation (between –30° and 30°) relates to associative memory retention (more or less forgetting across participants). Relationships between this phase locking measure and forgetting were not significant across all participants (*r* = .07, *p* = .642) or within either TMS condition (both *p*s > .493; Fig. S2B), suggesting that stronger coupling between reactivation and the peak GS phase is not reliably related to later memory. These results indicate that reactivation preferentially occurs around the peak of the ongoing GS fluctuation, a phase previously associated with arousal reductions (Liu et al., 2018), though the strength of this phase coupling was not reliably related to memory retention.

**Figure 4.**
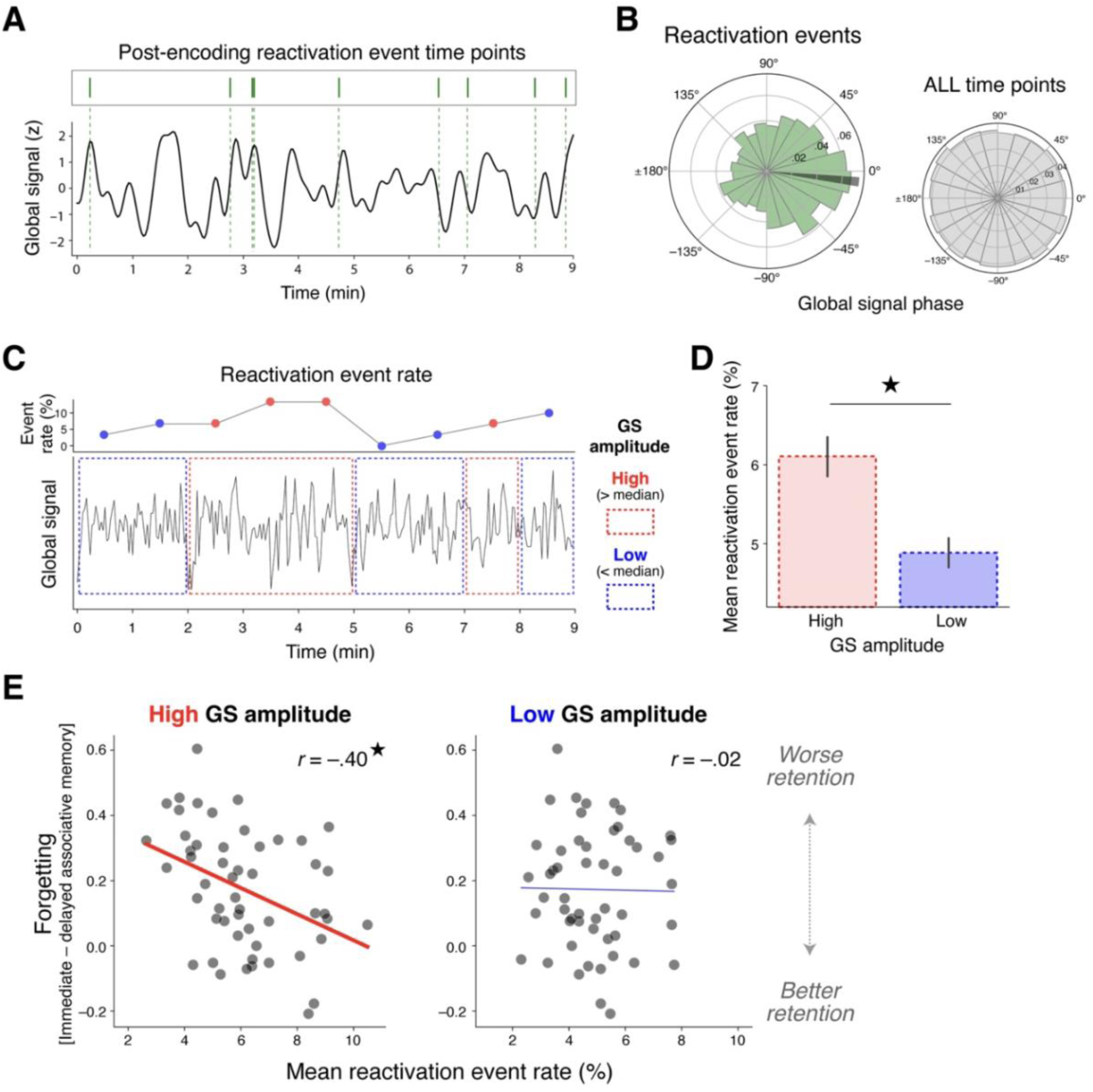
Coupling between the global signal (GS) dynamics and reactivation events and its relationship with memory. (A) An example GS time course (0.01-0.05 Hz band-pass filtered, z-scored), with reactivation event time points marked by green ticks and dashed vertical lines. (B) Distribution of GS phase during reactivation events (left) and all time points (right), aggregated across all participants. A phase of 0 degrees corresponds to the GS peak. (C) Example of the GS amplitude analysis. The GS time series is divided into successive 1-min windows, with windows classified as high (red) or low (blue) GS amplitude using a median split. The top line plot shows reactivation event rate in each successive time window. (D) Mean reactivation event rate in high-versus low-amplitude GS windows across participants (star indicates paired t-test, p < .001). (E) Relationships between mean reactivation event rate and associative forgetting (immediate - delayed associative memory accuracy), computed separately for high-GS (left) and low-GS (right) amplitude windows. Each data point represents one participant.

We next examined the relationship between reactivation and infra-slow fluctuations in the GS amplitude, as high GS amplitude has been linked with reduced arousal or vigilance across minutes (Liu & Falahpour, 2020; Wong et al., 2013). To do so, we divided each post-encoding rest scan into successive 1-min time windows and computed GS amplitude within each window (Gonzalez-Castillo et al., 2022), as shown for an example participant in Figure 4C. To test whether reactivation events are more prevalent during high versus low GS amplitude periods, we performed a median split on the 1-min windows based on each participant’s GS amplitude. Reactivation events occurred more frequently during high versus low GS amplitude time windows (*t_51_* = 4.02, *p* < .001) (Fig. 4D). The higher prevalence of reactivation during high GS amplitude windows did not significantly differ based on TMS site (Fig. S3A, mixed effects ANOVA: TMS site × GS window interaction: *F*_1,50_ = 2.70, *p* = .107, main effect of GS amplitude: *F*_1,50_ = 16.7, *p* < .001), further suggesting that, like GS phase coupling, reactivation-GS amplitude coupling reflects a stable rather than a malleable feature of reactivation dynamics. Consistent with the GS phase results above, these findings suggest that reactivation may be more prevalent during periods of reduced arousal, as reflected in slow, minute-long GS dynamics.

We next asked whether the relationship between reactivation and memory retention (forgetting) differs across high versus low GS amplitude windows. The correlation between reactivation rate and associative forgetting was significantly stronger during high compared to low GS amplitude windows (permutation test: *p* = .0348). Specifically, more reactivation during high GS amplitude windows was associated with better retention or reduced forgetting (*r* = –0.40, *p* = .003), while no relationship was found during low GS amplitude time windows (*r* = –0.02, *p* = .911). The relationship between reactivation during high GS amplitude windows and memory retention did not reliably differ between TMS sites (Fig. S3B; TMS × reactivation rate interaction: *b* = –.029 ± .027, *t* = –1.07, p = .290). These results indicate that the link between reactivation rate and memory retention is selectively observed during high GS amplitude periods, which may reflect reduced arousal (Wong et al., 2013), suggesting that reactivation may be more effective in supporting memory stabilization during lower arousal periods.

To further corroborate the above findings using a graded rather than a median-split measure of GS amplitude, we examined whether, within individual participants, GS amplitude correlated with reactivation event rates across time windows (*n* = 27) (see the inset of Fig. S3C for an example scatter plot). Consistent with the median-split approach, Spearman’s correlation between GS amplitude and reactivation rate was significantly above 0 across all participants (*t_51_* = 5.17, *p* < .001) (Fig. S3C) and did not differ between TMS groups (independent *t*-test: *t_50_*= 0.42, *p* = .679) (Fig. S3C). Similarly, stronger correlations between GS amplitude and reactivation rate were associated with less forgetting across participants (*r* = –0.30, *p* = .028), and this relationship did not differ between TMS groups (TMS × GS amplitude-reactivation Correlation interaction: *b* = .060 ± .219, *t* = .27, *p* = .786) (Fig. S3D). Together with the above results (Fig. 4E), these findings indicate that participants who exhibit better memory retention (reduced forgetting) show stronger coupling between reactivation and GS amplitude dynamics, potentially related to reduced arousal. Moreover, the preserved links between reactivation, GS dynamics, and their coupling with memory retention regardless of TMS site suggest that GS dynamics, previously linked to arousal, may be a relatively stable feature of awake reactivation.

### Brain States and Global Signal Dynamics Associated with LOC and Hippocampal Reactivation

To assess whether global brain dynamics linked to coordinated reactivation events reported above reflect properties specific to coordinated hippocampal-cortical co-reactivation, or whether they generalize to reactivation events identified in individual brain regions, we examined hippocampal and LOC reactivation separately (see Supplementary Results for full description). The two regions showed distinct coupling with whole-brain activity patterns and GS dynamics (Fig. S4A). LOC reactivation was preferentially coupled with the sensorimotor pole of the S-A axis (3A), while hippocampal reactivation was coupled with the association pole (3B). In addition, LOC reactivation was no longer coupled with the internally oriented state (brain state pole). Reactivation in the two regions also differed in the relationship with GS fluctuations (Fig. S4B): hippocampal events clustered at the GS peak, similar to coordinated reactivation events, whereas LOC events peaked earlier in the GS cycle.

## Discussion

Our findings show that spontaneous awake reactivation does not occur uniformly over time but is embedded within whole-brain dynamics, and its relationship to later memory depends on the ongoing brain state (see conceptual summary in Fig. 5). Reactivation was coupled with an internally oriented state characterized by DMN activation and DAN suppression, whole-brain activity patterns reinstated from stimulus encoding, and infra-slow GS dynamics previously associated with lower arousal. These relationships were behaviorally relevant: the expression of distinct whole-brain patterns was linked to opposing memory outcomes, and reactivation during high GS amplitude windows was linked to better retention. Brain stimulation selectively perturbed the coupling between reactivation and whole-brain activity patterns, while preserving coupling with GS dynamics.

**Figure 5.**
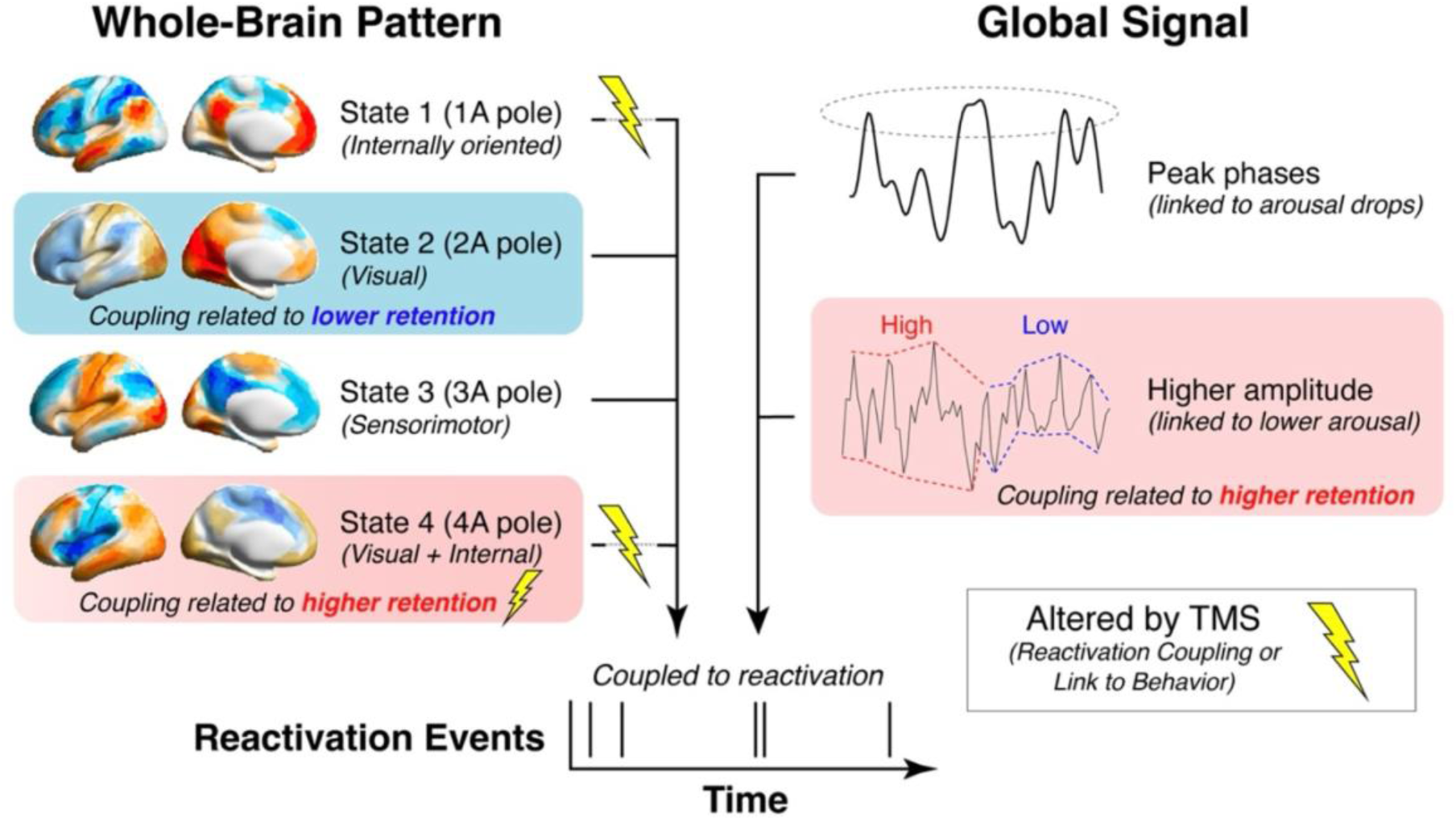
Summary of whole-brain activity patterns and global signal (GS) dynamics associated with memory reactivation. Schematic summary of global brain dynamics coupled with reactivation events. Reactivation preferentially occurred during an internally oriented brain state (State 1A), sensory states (States 2A and 3A), and a combined visual/internal state (State 4A) (left). Reactivation was also more likely to occur during peak phases and high-amplitude periods of the GS (right), both features putatively linked to lower arousal. Reactivation coupling with State 2A was related to poorer memory retention (blue), whereas coupling with State 4A and the global signal amplitude were related to better retention (red). TMS (lightning bolts) altered reactivation coupling with States 1A and 4A, and the link between State 4A and memory, but did not reliably alter coupling between reactivation and GS dynamics.

Spontaneous memory reactivation was coupled with an internally oriented brain state (1A brain state pole) characterized by DMN activation, DAN suppression, and associations with self-referential and autobiographical terms. This state was selectively expressed during resting reactivation but suppressed during matched online stimulus encoding events. This dissociation supports frameworks emphasizing opposing dynamics between external encoding and internal retrieval or consolidation (Buzsáki, 1989; Hasselmo & McGaughy, 2004; Honey et al., 2017) and mirrors empirical evidence for trade-offs between processing modes (Yang et al., 2025; Higgins et al., 2021; Duncan et al., 2012; Huijbers et al., 2011, 2012; Daselaar et al., 2009; Long & Kuhl, 2019). DMN engagement has also been linked to hippocampal ripples and reactivation (Higgins et al., 2021; Logothetis et al., 2012; Kaplan et al., 2016; Norman et al., 2019, 2021). Our findings connect this literature to slower fluctuations in fMRI brain state dynamics, raising the possibility that some identified large-scale activity patterns may reflect physiological states that support SWRs and reactivation.

The internally oriented activity pattern (1A) was coupled with both hippocampal and coordinated hippocampal-cortical reactivation, but not visual (LOC) reactivation, suggesting a role in hippocampal-cortical coordination beyond visual reinstatement alone. One possibility is that DAN suppression present in 1A may reduce competition from externally oriented attention and sensory processing, creating conditions favorable for hippocampal-cortical reactivation and internally generated memory representations. DMN engagement may promote hippocampal-cortical interactions, consistent with evidence of post-encoding reactivation and connectivity in DMN regions (Meyer et al., 2019; Yu et al., 2024). Given the DMN’s broad connectivity with the hippocampus and distributed cortical regions, DMN activation may provide a pathway for reactivation to propagate and recruit more distributed cortical representations (Kaefer et al., 2022; Joshi et al., 2026; Margulies et al., 2016). Our findings place these proposed DMN-facilitated interactions within a broader network context of concurrent DMN activation and DAN suppression.

Whereas the internally oriented 1A pattern was expressed during resting reactivation and suppressed during encoding, States 2-4 showed congruent expression across both reactivation and encoding. States 2-4 also shared prominent loading of the visual network, largely absent in State 1, consistent with reinstatement of sensory encoding-related processing. Although reinstatement of these brain states parallels reactivation itself, the two measures capture different spatial scales (Coutanche, 2013): reactivation reflects multi-voxel patterns within regions, while brain states reflect whole-brain patterns of regionally averaged activity, defined from baseline rest. Together, spontaneous reactivation occurs during a combination of brain states that reflect reinstated stimulus-processing patterns and internally oriented activity.

Although reactivation was coupled with all four brain states, only a subset was related to memory retention. Expression of the internally oriented state (1A) during reactivation was not reliably linked to memory retention, in contrast to States 2 and 4. While the null association for State 1 does not rule out a smaller effect, this pattern suggests that brain states permissive for reactivation may differ from those that support memory stabilization. Notably, both 1A and 4A featured DMN activation, but only 4A expression was reliably associated with retention, suggesting that DMN engagement may not be sufficient for memory stabilization, which may instead depend on the broader network configuration during reactivation. Beyond DMN activation, the 4A pattern included co-activation of limbic (temporopolar and orbitofrontal cortex) and visual networks, along with suppression of the ventral attention and somatomotor networks. Consistent with the reinstatement of encoded visual content, this state was also associated with terms related to visual episodic memory (’fusiform’, ‘faces’, ‘episodic’).

The broad co-activation of visual and memory-related networks in 4A may represent a favorable state for visual memory stabilization, coordinating sensory (visual) areas and those supporting integration, schematic representations, and semantic knowledge (Gilboa & Marlatte, 2017; Ritchey & Cooper, 2020; van Kesteren et al., 2012; Lambon Ralph et al., 2017; McClelland et al., 1995; Squire et al., 2015; Klinzing et al., 2019). This pattern is aligned with prior work linking post-encoding functional connectivity over minutes in similar regions to subsequent behavior (Tambini et al., 2010; Schlichting & Preston, 2014, 2016; Murty et al., 2017; Hermans et al., 2017; Meyer et al., 2019; Tompary & Davachi, 2024). Our findings show that memory retention is linked to coordinated activation of large-scale visual and memory-related networks at the moments when spontaneous reactivation occurs. Additionally, the suppression of the ventral attention and somatomotor networks in 4A may help shield reactivated memory representations from competing processing, consistent with the role of ventral attention/salience systems in reorienting attention toward behaviorally relevant events (Seeley et al., 2007; Menon & Uddin, 2010), recent proposals that this network supports externally oriented, action-related processing (Dosenbach et al., 2025), and evidence that declarative and procedural memory systems compete for offline processing (Robertson et al., 2004; Brown & Robertson, 2007). Together, coordinated activation across visual and memory-related networks with reduced competing processing may provide a favorable whole-brain context for consolidation.

Unlike the 4A brain state pole, whose broader network configuration was associated with better retention, 2A strength during reactivation was associated with worse retention. 2A strongly loaded onto the visual network and was associated with lower-level visual terms (‘visual cortex’, ‘occipital’, ‘primary visual’). This pattern may reflect a sensory-dominated reactivation format without the broader co-activation of memory-related networks observed in 4A. Another possibility is that this brain state may index sensory attention or a more externally oriented processing mode that could interfere with concurrently reactivated representations (Dewar et al., 2012, 2014; Risius et al., 2019). Regardless of the particular mechanism, the opposing relationships of 2A and 4A with memory suggest that the broader whole-brain configuration accompanying awake reactivation may shape whether reactivation benefits or impairs memory.

Beyond whole-brain activity patterns, reactivation clustered near the peak of the global fMRI signal (GS), linking hippocampal-cortical reactivation to periods of globally elevated BOLD signal. This finding mirrors observations that hippocampal ripples and replay are coupled to widespread whole-brain activity and large-scale neural cascades (Yang et al., 2024; Liu et al., 2022; Logothetis et al., 2012; Nitzan et al., 2022). Reactivation was also more frequent during high GS amplitude windows, when its occurrence was associated with memory retention. Both the GS peak and higher GS amplitude have been associated with reduced arousal (Bolt et al., 2025; Liu et al., 2018; Gonzalez-Castillo et al., 2022; Wong et al., 2013), raising the possibility that lower arousal periods, as indicated by GS fluctuations, may represent favorable windows for memory stabilization. These periods may also coincide with internally oriented network configurations, reflecting a broader shift away from externally oriented processing to a state permissive of spontaneous reactivation. However, the GS is not a specific measure of arousal and reflects multiple neural and physiological processes, including respiratory and cardiovascular contributions (Bolt et al., 2025). Simultaneous EEG-fMRI, pupillometry, and autonomic recordings are needed to test whether these windows correspond to lower arousal states.

Perturbing post-encoding brain activity via TMS modified the coupling between reactivation and whole-brain activity patterns, as well as the relationship between State 4 expression and memory retention. These findings suggest that the large-scale network context expressed during reactivation, and its relationship to memory, are sensitive to brain stimulation. This is consistent with theta-burst TMS altering large-scale network activity by influencing regions connected to the stimulation site (Bestmann & Feredoes, 2013; Bergmann & Hartwigsen, 2021; Gratton et al., 2013; Hebscher & Voss, 2020). In contrast, reactivation remained coupled with GS features previously associated with arousal, suggesting that whole-brain activity patterns and arousal-related GS fluctuations may be partially dissociable aspects of the ongoing brain state.

Our single region results also offer insight into the large-scale organization of spontaneous reactivation. Recent work proposes that hippocampal reactivation propagates along the principal gradient (Margulies et al., 2016), or the sensorimotor-association (S-A) axis, through transmodal association cortex toward unimodal sensory regions (Kaefer et al., 2022). Consistent with this proposed organization, we found that hippocampal reactivation was coupled with the association pole of the S-A axis (3B), whereas LOC reactivation was coupled with its sensorimotor pole (3A). However, coordinated hippocampal–LOC events were also linked to the sensorimotor pole, and LOC reactivation peaked approximately 45° earlier in the infra-slow GS cycle than hippocampal reactivation. Although this phase offset indicates different temporal organization of hippocampal and cortical reactivation relative to the GS, it does not establish that sensory reactivation precedes hippocampal reactivation. This phase offset may reflect infra-slow fluctuations that bias when reactivation occurs in different regions, consistent with infra-slow arousal dynamics organizing brain-wide activity during wakefulness (Raut et al., 2021; Yang et al., 2026) and memory-related processes during sleep (Jacobsen et al., 2026; Dimitriades et al., 2026).

We found that even within eyes-closed rest, often treated as a relatively homogeneous ‘offline’ state, spontaneous reactivation and its relationship to memory depended on the ongoing brain state. While behavioral studies typically compare the memory benefits of post-learning rest periods with externally oriented tasks (Brokaw et al., 2016; Craig et al., 2015; Dewar et al., 2014), the current results suggest that opportunities for consolidation fluctuate within rest itself. This interpretation converges with other work highlighting that dynamic variability in SWR rates track arousal-related pupil fluctuations (McGinley et al., 2015; Yang et al., 2025; see also Chang et al., 2025) and that distinct offline waking states can be isolated using EEG, pupillometry, and behavior (Wamsley & Summer, 2020; Wamsley et al., 2023; Wamsley & Collins, 2024). These approaches converge to highlight heterogeneity within awake periods; here, our findings link ongoing dynamics to the timing of individual hippocampal-cortical reactivation events and their relationship to later memory. Future simultaneous EEG-fMRI with pupillometry could test whether physiologically and behaviorally defined offline states coincide with brain states identified here and whether reactivation is most beneficial when distinct measures converge.

Fluctuations in awake brain states favorable for reactivation raise the question of how such states relate to memory processing during sleep. Quiet wakefulness and NREM sleep share features including SWRs, reactivation, and relatively reduced cholinergic tone (Hasselmo, 1999; Hasselmo & McGaughy, 2004), and behavioral work suggests partially overlapping memory benefits (Tucker et al., 2020; Wamsley, 2019). Within NREM sleep, infra-slow noradrenergic and autonomic fluctuations may organize periods of spindle expression and associated memory outcomes (Jacobsen et al., 2026), while respiration modulates sleep oscillations and their coupling with memory reactivation (Schreiner et al., 2023). Together with our findings, this work suggests that consolidation fluctuates within both NREM sleep and awake rest, with particular recurring brain and physiological states favoring reactivation and consolidation. However, the physiological properties of these states, as well as the nature and function of reactivation, may likely differ between wakefulness and sleep (Roumis & Frank, 2015).

A recurring puzzle is that awake reactivation is not uniformly related to memory benefits. Studies of exogenously cued awake reactivation, or targeted memory reactivation (TMR), have reported diverging outcomes (i.e., memory benefits, impairments, or null effects) (Rudoy et al., 2009; Diekelmann et al., 2011; Oudiette et al., 2013; Schreiner & Rasch, 2015; Tambini et al., 2017; Alm et al., 2019; Hoffman et al., 2024), with outcomes also depending on conscious processing (Tal et al., 2024). Similarly, studies of spontaneous consolidation during awake rest have sometimes reported variable memory benefits (Martini et al., 2018, 2020; Varma et al., 2017; Humiston et al., 2019; Tanriverdi et al., 2023). Our findings suggest that the ongoing brain state during reactivation may help explain this variability: distinct whole-brain activity patterns expressed during reactivation were associated with opposing memory outcomes, and the relationship between reactivation and retention differed depending on the ongoing GS amplitude. These findings raise the possibility that awake reactivation, whether cued or spontaneous, may stabilize memories in some brain states but destabilize or increase their susceptibility to interference in others. An intriguing question is whether these brain states also differ in conscious access to reactivated content, which may potentially relate to the differing consequences of conscious and unconscious cueing (Tal et al., 2024). More broadly, these results and others (Wamsley & Summer, 2020; Wamsley et al., 2023) suggest that the ongoing brain state may shape the outcome of awake reactivation manipulations.

In summary, our findings show that spontaneous awake reactivation and its relationship to memory are state-dependent, even within eyes-closed rest. More broadly, our findings are consistent with emerging frameworks linking hippocampal activity and cognition to large-scale brain dynamics and physiological state (O’Callaghan et al., 2021; Colcombe, 2025). Together, these results suggest that fluctuations in the ongoing brain state may create favorable windows for awake memory consolidation, shaping both when reactivation occurs and whether it is associated with memory stabilization or forgetting.

## Methods

### Participants

The data presented here have been reported in a previous study (Tambini & D’Esposito, 2020). Data were collected from 62 human participants, and after exclusion, 52 participants remained for analyses (28 females and 24 males; aged 18-26 with a mean of 20.6 years). See Tambini and D’Esposito (2020) for a detailed description of participant exclusion criteria, which included high head motion and chance-level immediate memory performance. All participants provided prior informed consent in accordance with the Committee for Protection of Human Subjects at the University of California, Berkeley.

### Experimental Procedures

#### Overview

Participants completed two experimental visits: a baseline session followed by a main session. The baseline session, described below, included MRI scans necessary for TMS procedures performed during the main session, as well as high-level visual category functional localizer scans. The main session consisted of the following procedures, in order (Fig. 1A): a baseline rest fMRI scan (9 min), five incidental visual encoding scans (9.6 min each), an immediate memory test (outside the MRI), TMS (outside the MRI), post-encoding rest scans (three 9 min BOLD scans interleaved with 5 min Arterial Spin Labeling [ASL] scans), and delayed memory tests (outside the MRI, administered 3 hrs later and next day). Participants were not informed about the memory tests until immediately before each test. Memory was measured both immediately after encoding and at delayed time points to obtain a behavioral index of consolidation via the change in memory over time (immediate compared to delayed memory). TMS was applied before post-encoding rest, with its influence expected to span the duration of the ∼40 min post-encoding rest period (Wischnewski & Schutter, 2015). The TMS manipulation was originally designed to test the causal role of post-encoding activity in memory consolidation, the primary goal of this dataset (see Tambini & D’Esposito, 2020); here, we test the influence of TMS on memory reactivation dynamics. Reactivation dynamics were examined during the three post-encoding BOLD fMRI scans (27 min total). ASL data are not reported here because of their substantially lower spatial and temporal resolution compared to BOLD fMRI. Notably, the 27 min post-encoding rest period provided unusually extended sampling of post-encoding reactivation dynamics compared to most studies, which typically collect 5-10 min of post-encoding fMRI data (e.g., Tambini & Davachi, 2013; Staresina et al., 2013; Schapiro et al., 2018; Tanriverdi et al., 2023; Yu et al., 2024).

#### Rest

During pre- and post-encoding rest scans (each lasting 9 min), participants were instructed to rest, think about anything that naturally comes to mind, and close their eyes while remaining awake.

#### Encoding task

Participants incidentally encoded object-face pairs while performing an imagery task. During encoding, participants viewed 180 object-face pairs across five scans, with each stimulus presented to the left or right of a central cross. Each encoding scan (lasting 9.6 min) contained 36 trials. Each trial consisted of a brief central cross (0.25 s), followed by an object-face pair (5.75 s), a vividness rating (1 s), and an active baseline task (9 s). Participants were instructed to vividly imagine the person pictured interacting with the object and rate the vividness of their mental imagery during the 1s rating period (participants were not explicitly instructed to encode images in preparation for memory tests). The active baseline task involved indicating the direction of an arrow (left or right) via self-paced button responses, designed to reduce hippocampal activity during inter-stimulus intervals (Stark & Squire, 2001). The visual stimuli presented during encoding scans and memory tests consisted of 244 object-face pairs. See Tambini and D’Esposito (2020) for more details regarding the stimuli.

#### Memory testing

Memory tests were designed to assess memory for single items as well as object-face associations. Here we focus on associative memory, rather than item memory, as it is more closely related to hippocampal function and post-encoding consolidation indices in prior work studying spontaneous memory reactivation and hippocampal-cortical interactions (Tambini et al., 2010; Gruber et al., 2016; Murty et al., 2017). During memory tests, participants viewed object-face pairs and judged each pair as ‘Intact’ (seen together during encoding), ‘Rearranged’ (seen during encoding, but paired with other stimuli), or ‘New’ (one or both items not seen during encoding). Upon choosing ‘New’, participants were prompted to indicate which item was new: object, face, or both items. Participants were asked to be as accurate as possible with no time constraint. A separate subset of the encoded stimuli was tested during each memory test. The immediate test was administered just after encoding, and two delayed tests were administered 3 and 24 hours after TMS. Given prior evidence from this dataset linking the reactivation measures examined here primarily to same-day memory retention, we focus on the same-day (3 hr) delayed memory test (Tambini & D’Esposito, 2020). The immediate test consisted of 60 object-face pairs (32 intact, 8 rearranged, 8 one new/one old, 12 both new), and each delayed test contained 92 object-face pairs (48 intact, 12 rearranged, 16 one new/one old, 16 both new).

#### TMS Procedures

To test the causal role of post-encoding brain activity in memory consolidation (the primary goal of our prior work, Tambini & D’Esposito, 2020), TMS was applied shortly after the immediate memory test to LOC or a Control site (primary somatosensory cortex) using a standard continuous theta-burst TMS protocol (triplets of 50 Hz pulses, administered at 5 Hz for 40 s; Huang et al., 2005). Participants were randomly assigned to one of the two TMS sites (LOC or the Control site). The LOC TMS site was defined as the site of maximal responsiveness to objects (objects > scrambled objects contrast from the functional localizer scans) in right lateral occipital cortex (as in prior TMS work, Pitcher et al., 2011). Right LOC (as opposed to left LOC) was targeted as prior work found post-encoding hippocampal-LOC interactions that were greater for right versus left LOC (Tambini et al., 2010) and other studies have successfully interfered with object processing by targeting right LOC (Pitcher et al., 2011). The Control TMS site (dorsomedial somatosensory cortex, S1) was defined anatomically (the most superior portion of the right postcentral gyrus that fell within a gray matter mask). The TMS sites for all participants are shown in MNI space in Figure 1B of the original study (Tambini & D’Esposito, 2020). An active control condition was used to ensure that any influence of TMS was not due to non-specific effects of stimulation or potential changes induced by the acoustic or tactile scalp sensations associated with TMS. As in prior work (Gratton et al., 2013; Nee & D’Esposito, 2017; Tambini et al., 2018), dorsomedial S1 was chosen as the active control site since it demonstrates relatively local connectivity within somatosensory cortex (Yeo et al., 2011) and corresponds to the leg somatosensory representation, making it unlikely that S1 cTBS would influence hippocampal function related to memory consolidation. Additional details regarding TMS procedures are described in Tambini and D’Esposito (2020).

#### Baseline session

Functional localizer as well as anatomical (T1) scans were acquired during a baseline session (before the main session). The localizer data were used to identify the LOC region of interest (ROI) as well as to obtain category-specific activity patterns associated with face and object processing. Standard high-level visual functional localizer scans were collected, during which participants viewed stimulus categories in a blocked fashion (objects, faces, scenes, and phase scrambled images, using distinct stimuli from the main session) while performing a one-back task (i.e., pressed a button when the same image was repeated). Each scan consisted of six blocks of each stimulus category (24 blocks in total) interleaved with seven blocks of passive fixation. Each block lasted 16 s and contained 20 stimuli (0.3 s stimulus presentation followed by 0.5 s blank). Two 8.27 min localizer scans were collected for each participant.

### Behavioral Data Analysis

To assess the functional relevance of brain state dynamics during memory reactivation, we focused on memory retention as a behavioral index of memory consolidation (i.e., a forgetting score, as in prior work, e.g., Brokaw et al., 2016; Wamsley & Summer, 2020). We focused on associative memory specifically, given its link with hippocampal function and post-encoding consolidation mechanisms (Davachi, 2006; Mayes et al., 2007; Tambini et al., 2010; Gruber et al., 2016; Murty et al., 2017). Associative memory accuracy was measured as the associative hit rate (i.e., proportion of intact trials labeled as intact) minus the associative false alarm rate (i.e., sum of the proportion of rearranged and new trials labeled as intact). Forgetting, inversely related to retention, was calculated as immediate minus delayed associative memory accuracy (i.e., a higher score indicating more forgetting or poorer retention).

### Functional MRI Data Acquisition and Preprocessing

Scanning was performed using a 3T Siemens TIM/Trio MRI system with a 12-channel whole-head coil at the UC Berkeley Henry H. Wheeler Jr. Brain Imaging Center. High-resolution T1-weighted (magnetization-prepared rapid-acquisition gradient echo) images were acquired during the baseline session (voxel size = 1 × 1 × 1 mm; TR = 2300 ms; TE = 2.98 ms; flip angle = 9; matrix = 240 × 256 × 160). Functional T2*-weighed images were collected using a gradient-echo planar pulse imaging (EPI) sequence (voxel size = 3 × 3 × 3 mm; TR = 2 s; TE = 28 ms; field of view = 210 mm; 32 slices, 0.6 mm interslice gap, flip angle = 78°). The EPI scans contained 270 (rest), 288 (encoding), and 248 (functional localizer) volumes (TRs), after discarding the first five volumes to allow for T1 equilibration.

The imaging data were preprocessed using SPM8 as described previously (Tambini & D’Esposito, 2020). Functional data were first corrected for slice timing differences, followed by motion correction and low-frequency trend removal (< 0.009 Hz, using spm_filter). Each participant’s functional data were co-registered to their T1-weighted anatomical image using the mean functional image from each session. Functional data were spatially smoothed using a 6-mm full-width at half-maximum (FWHM) isotropic Gaussian kernel, except for the hippocampal pattern reactivation analysis (as in Tambini & Davachi 2013, Tambini & D’Esposito, 2020). All analyses were performed in each participant’s native space.

Denoising of fMRI data was performed via nuisance signal regression and excluding motion-contaminated volumes from all analyses (motion scrubbing; Power et al., 2012). Specific nuisance signals regressed out of all fMRI runs included: aCompCor regressors (the top principal components, PCs, from WM and CSF masks, Behzadi et al., 2007), six motion parameters, and the 1st temporal derivatives of the six motion parameters. Nuisance signal regression was performed in a voxel-wise manner for each fMRI scan (for a full description see Tambini & D’Esposito, 2020). Brain state analyses were performed on low-pass filtered data (< 0.1 Hz, 4th order Butterworth filter applied after nuisance signal removal). Lastly, motion scrubbing was performed for all scans by discarding volumes or time points surrounding head motion events from all analyses (one volume before and two volumes after all events of framewise displacement > 0.5 mm; Power et al., 2012; Balachandrasekaran et al., 2022).

### LOC and Hippocampal Reactivation Analysis

To examine how reactivation dynamics are related to global brain states, we identified reactivation events that occurred jointly across the hippocampus and a cortical site involved in processing encoded stimuli, namely LOC. This region was specifically chosen based on prior work showing reactivation in or memory-related hippocampal interactions with similar or nearby portions of occipito-temporal cortex during post-encoding rest (Tambini et al., 2010; Deuker et al., 2013; Schlichting & Preston, 2014; de Voogd et al., 2016; Murty et al., 2017; Kark & Kensinger, 2019). We took this approach rather than examining reactivation in a single brain region for two reasons. First, joint or coordinated activity between the hippocampus and cortex is thought to mechanistically facilitate memory stabilization across hippocampal-cortical networks, representing an important feature of how memory representations evolve over time (McClelland et al., 1995; Maingret et al., 2016; Geva-Sagiv et al., 2023; Tompary & Davachi, 2017). Second, these two regions are affiliated with distinct large-scale networks, allowing us to avoid biasing brain state activity linked with reactivation towards the activation of any particular network. For example, hippocampal reactivation events may be expected to co-vary with a brain state that strongly weights the DMN, as hippocampal activity tends to co-vary with DMN activity (Vincent et al., 2006). However, joint reactivation events across the hippocampus and LOC would not be expected to show the same degree of bias towards a single large-scale network, since LOC tends to be affiliated with the ‘visual’ network (Yeo et al., 2011).

To isolate reactivation events, LOC was functionally defined in each participant as object-responsive portions of lateral occipital cortex and posterior fusiform cortex using standard criteria (greater activation for objects vs. scrambled images in functional localizer scans, p < .0001 uncorrected). The details of general linear models (GLMs) used to analyze the functional localizers scans are reported in Tambini and D’Esposito (2020). Briefly, standard GLMs were constructed that contained regressors for each stimulus category block (faces, objects, scenes, and scrambled images). To measure the reactivation of object and face stimuli in LOC, we used the same approach as previously reported (Tambini & D’Esposito, 2020). Specifically, we obtained two separate template multi-voxel patterns (face and object) from the localizer scans for each participant. Each template pattern (face, object) was constructed by extracting the time series for all LOC voxels from each localizer scan, z-scoring the BOLD signal over time, and calculating the average value (z-scored signal) during the approximate peak of the hemodynamic response for each voxel (4-8^th^ TRs or 6-14 s from the onset of face and object category blocks), resulting in a multi-voxel activity pattern. To measure the similarity of spontaneous resting activity patterns to face or object template patterns, we extracted the time series of LOC voxels during each rest scan and z-scored the BOLD signal in each voxel over time. We then correlated each of the two template multi-voxel patterns (face or object patterns derived from the localizer scans) with the activity pattern at each time point (i.e., TR) during rest (Fig. 1C). The resulting Pearson’s correlation coefficients were Fisher’s z-transformed. We used a thresholding approach to identify reactivation events from the similarity (i.e., correlation) dynamics, as in previous studies (Staresina et al., 2013; Schapiro et al., 2018; Tanriverdi et al., 2023; Yu et al., 2024). For each participant, LOC reactivation events were defined as rest volumes (time points) with high similarity or correlation to either the object or face template patterns (> 1.5 standard deviations, SD, above the mean z-transformed correlation across baseline and post-encoding rest volumes, Fig. 1D). This threshold of 1.5 SD has been used in prior work (Staresina et al., 2013; Schapiro et al., 2018; Yu et al., 2024).

To isolate hippocampal reactivation events, each participant’s hippocampal ROI was anatomically defined using FSL’s FIRST (Patenaude et al., 2011), visually inspected, and manually corrected when necessary to conform to anatomical boundaries described by Pruessner et al. (2000). As previously described (Tambini & D’Esposito, 2020), voxels with low signal (< 2 SD below the whole brain mean) or high variance (> 4 SD above the median variance across hippocampal voxels) in any encoding or rest scans were excluded from the hippocampal ROI in each participant. Since hippocampal activity patterns are typically not sensitive to stimulus categories (Huffman & Stark, 2014), we examined hippocampal reactivation using template patterns derived from the encoding data. Following prior work using the same or similar experimental designs (Tambini & Davachi, 2013; Tambini & D’Esposito, 2020), we isolated hippocampal template encoding patterns by performing principal component analysis (PCA) on unsmoothed hippocampal ROI voxel data. Specifically, the time series of hippocampal voxels were extracted and z-scored over time in each encoding scan, concatenated across all encoding scans for each participant, and PCA was performed. We focused on the top three encoding PCs, or patterns that explained the largest amount of variance in the hippocampal encoding data for subsequent reactivation analysis. The top three PCs were chosen based on prior work showing that: 1) the top three PCs capture reliable stimulus-evoked activity during encoding and 2) reliable reactivation evidence was observed for the top three hippocampal encoding PCs in a similar dataset (Tambini & Davachi, 2013). To measure hippocampal reactivation, we extracted the time series of hippocampal voxels during each rest scan and z-scored the BOLD signal in each voxel over time. We then calculated the similarity between each encoding pattern (PC) and the hippocampal pattern at every time point during rest using a component score, obtained by projecting the voxel weights of each PC onto the hippocampal activity pattern at each time point during rest (Fig. 1C). Hippocampal reactivation events were identified using the same threshold as LOC reactivation (timepoints > 1.5 SDs above the mean component score for any of the top three PCs; Fig. 1D).

After identifying reactivation events separately in the hippocampus and LOC, we isolated coordinated reactivation events. Coordinated events were defined as time points of hippocampal reactivation in which LOC reactivation concurrently occurred at the same or an adjacent time point (Tanriverdi et al., 2023; Fig. 1D) and were used for all further analyses except where noted.

### Brain State Identification

To measure dynamic fluctuations in brain states, we characterized moment-to-moment spatial patterns of whole-brain activation. As in prior work, we used a data-driven approach to identify prominent spatial activation patterns by performing k-means clustering of fMRI activation patterns (Liu et al., 2013; 2018). This approach identifies a specified number (*k*) of clusters, or recurring spatial activity patterns, that capture substantial variance in the data, such that similar spatial activity patterns expressed at non-adjacent timepoints are assigned to the same activity pattern or cluster number (1 to k). We set the number of clusters (i.e., spatial activity patterns) to *k* = 8, which recent work identified as a particularly reliable number for resting state fMRI dynamics in humans (Gutierrez-Barragan et al., 2024). To avoid biasing the definition of whole-brain activity patterns by the reactivation of stimuli encoded in our experiment, we identified brain states using the baseline (pre-encoding) rest data. K-means clustering was performed on ROI-level time series, both to reduce the computational burden (Gutierrez-Barragan et al., 2024; Lee et al., 2024) and to characterize large-scale, regional activity patterns rather than within-region multivoxel patterns used to identify reactivation events. Specifically, we used 1,000 cortical and 54 subcortical ROIs from the Schaefer and Tian atlases, respectively (Schaefer et al., 2018; Tian et al., 2020). MNI-defined ROIs (1,054) were first transformed into study-specific template space (derived using Advanced Normalized Tools; Avants et al., 2011) and then into each participant’s native space. The time series of each ROI was extracted from each baseline rest scan, z-scored over time, concatenated across participants, and z-scored as a final step. K-means clustering was performed on the concatenated data (1054 ROIs x timepoints across all participants) using Matlab.

To visualize the spatial activation pattern of each k-means cluster, we generated a spatial map by averaging the activity in each ROI across baseline rest volumes or time points labeled as that cluster (Gutierrez-Barragan et al., 2019; 2024; Fig. 2A, shown on cortical surface representations). Consistent with previous work (Gutierrez-Barragan et al., 2024; Thompson et al., 2014), the resulting spatial maps formed pairs that were highly anti-correlated in both their spatial patterns and temporal expression (see Figs. 2A and 3A). We therefore conceptualized each pair as a brain state with two opposing poles or spatial activation patterns, labeled ‘A’ and ‘B’ (e.g., State 1 comprises the 1A and 1B poles). Thus, throughout the manuscript, we refer to the eight k-means-derived spatial patterns as the opposing A and B poles of four brain states. Although individual k-means clusters are sometimes referred to as singular brain states in the literature (e.g., Gutierrez-Barragan et al., 2019; Maltbie et al., 2022; Cornblath et al., 2020), we use the term ‘state’ here to refer to the paired A/B configuration to facilitate interpretation of the opposing spatial patterns and their continuous expression along a shared axis.

To further facilitate interpretation of the four identified brain states, we characterized the large-scale network activation profile of each brain state pole (e.g., 1A, 1B) by calculating the mean activation value (i.e., loading) within each of seven canonical large-scale cortical networks (Yeo et al., 2011; Fig. 2A, see bar plots below each map). In addition, we identified terms associated with the spatial pattern of each pole in the neuroscience literature, extracted from a Neurosynth meta-analysis (Yarkoni et al., 2011). Specifically, after removing duplicates and combining highly overlapping terms (e.g., ‘default’ and ‘DMN’), we selected the 10 terms most positively and 10 terms most negatively associated with each spatial map and visualized them in word clouds with text size reflecting the absolute correlation values (Fig. 2A). Since the A and B poles of each brain state are strongly anticorrelated in their spatial patterns, and thus yield highly similar (but inverted) associations, for simplicity we show word clouds only for the ‘A’ pole.

### Post-Encoding Brain State Analysis

To determine which brain states were associated with post-encoding memory reactivation, we first classified each post-encoding rest volume or time point as belonging to one of the eight spatial activity patterns corresponding to the A and B poles of the four brain states. In a secondary approach (described below), we also calculated the graded strength of each brain state. To do so, we first extracted the time series of each of the 1054 ROIs in each post-encoding rest scan and z-scored each time series over time. We then calculated the similarity (Pearson’s correlation coefficient) between the activity pattern of each post-encoding rest time point and each of the eight spatial activation maps (across 1054 ROIs), yielding eight correlation coefficients per time point as continuous indices of similarity to each whole-brain activity pattern. Each post-encoding rest time point was then assigned, in a winner-take-all fashion, to the spatial activity pattern (A or B pole of each brain state) with the highest similarity (i.e., correlation).

To test whether coordinated reactivation is linked with specific brain states, we quantified how often each of the eight spatial activity patterns, corresponding to the A and B poles of the four brain states, occurred during reactivation events (i.e., occurrence rate). The occurrence rate of each state pole (i.e., 1A, 1B, etc.) during reactivation was calculated as the proportion of reactivation time points labeled as each pole. To examine whether brain state expression was specifically associated with reactivation events, we also calculated occurrence rates of each brain state pole across all time points during post-encoding rest. We then derived normalized occurrence rates by dividing occurrence rates during reactivation events by the occurrence rates across all time points; a rate higher (or lower) than 1 indicates reactivation occurs more (or less) frequently than expected based on overall prevalence. To statistically test whether specific brain state poles preferentially occur during reactivation, we performed a repeated-measures ANOVA on the normalized occurrence rates with a factor of brain state pole, followed by one-sample t-tests (against 1) for each A and B pole. The resulting p-values were corrected for multiple comparisons using a Benjamini-Hochberg false discovery rate (FDR).

To test whether the brain activity patterns prominent during reactivation are specific to offline rest or are also engaged during encoding, we performed a parallel analysis of the encoding data. Specifically, we identified ‘encoding’ events from encoding scans — analogous to reactivation events during rest — by isolating time points with high similarity to the template activity patterns in the hippocampus and LOC (jointly across regions). We then classified each encoding time point into one of the eight spatial activity patterns using the same procedure as the rest scans, and calculated the normalized occurrence rates of each state pole during encoding events that fell within stimulus processing periods (i.e., 4-12 s after stimulus onset on each trial). To test whether brain state pole prevalence differs between reactivation and encoding events, we conducted a two-way repeated-measures ANOVA on the normalized rates, with brain state pole (eight levels) and time period (encoding vs. rest) as factors. We then performed one-sample t-tests (against 1) on the normalized rates during encoding events, as well as pairwise comparisons between reactivation and encoding normalized rates for each brain state pole (all Benjamini-Hochberg FDR-corrected).

As a complementary approach to categorical brain state labels (i.e., labeling each time point as a single brain state pole), we quantified the graded strength of each brain state to capture continuous variation in state expression along each brain state axis. Since the A and B poles of each brain state were strongly anticorrelated, we combined them into a single continuous measure for each of the four brain states (see Gutierrez-Barragan et al., 2024, for the same approach). Specifically, we defined graded brain-state strength by treating similarity to the A pole as positive and similarity to the B pole as negative, and averaging the two values (i.e., correlation-with-A minus correlation-with-B divided by two; Fig. 3A). Thus, positive strength values indicate greater expression of the A pole, whereas negative values reflect greater expression of the B pole. To quantify brain state strength during reactivation, we calculated the mean state strength during reactivation events and subtracted the mean state strength across all post-encoding time points to account for any non-reactivation-specific bias in state expression, such that positive or negative values indicate shifts toward the A or B poles, respectively. Reactivation-related brain state strength values were compared to 0 using one-sample t-tests with FDR correction across the four brain states. In addition, to test whether TMS influenced brain state strength during reactivation, we performed a mixed ANOVA on the mean state strength value (reactivation minus all time points) with factors of TMS site (LOC vs. Control) and brain state (four states). To isolate which brain states differed between TMS sites, pairwise independent t-tests were performed between the two TMS groups for each brain state.

To examine whether brain state expression during reactivation is related to memory retention, we focused on graded brain state strength, as it captures continuous variation in state expression beyond categorical classification or occurrence measures. To do so, we measured associative memory retention via forgetting (i.e., immediate minus delayed associative memory accuracy). We first tested the relationship between memory retention and brain state strength during reactivation across all participants via multiple regression, with the four brain state strength measures (reactivation minus all time points) entered simultaneously as predictors of associative forgetting. This multiple regression approach allowed us to estimate unique relationships between each brain state strength during reactivation and memory, while accounting for shared variance across the four brain states. While our primary analysis identified brain state-memory relationships across all participants, regardless of TMS site, we also tested whether these relationships were modulated by TMS. Specifically, we ran a separate regression for each brain state with three predictors of associative forgetting: brain state strength during reactivation, TMS group (categorical factor, mean centered), and their interaction term. Finally, to characterize these relationships within each TMS condition, we measured correlations between brain state strength during reactivation and associative forgetting separately in each TMS condition.

### Global Signal (GS) Analysis

We examined reactivation events in relation to slow fluctuations of the GS, which has been related to slow arousal changes in prior work (Gonzalez-Castillo et al., 2022; Liu & Falahpour, 2020). The GS was calculated by averaging the BOLD time-series across all voxels within each participant’s brain mask. We assessed the GS phase in the infra-slow frequency range, previously related to physiological arousal fluctuations (Bolt et al. 2025; Liu et al., 2018; Raut et al., 2021). To do so, the time course of the GS for each participant and each post-encoding rest scan was band-pass filtered from 0.01 to 0.05 Hz (Raut et al., 2021), using a Butterworth filter, and then z-scored (Fig. 4A). We then computed the instantaneous phase of the filtered GS using the Hilbert transformation.

To assess whether reactivation occurs more frequently during specific GS phases, we isolated the GS phase at each reactivation event and aggregated the phase values across all participants into a single distribution (Fig. 4B). We then statistically tested whether the distribution differs from circular uniformity using a Rayleigh test. To investigate whether the strength of coupling between reactivation and GS phase is behaviorally relevant, we used a data-driven approach to first identify the GS phase around which reactivation events clustered and then test whether variation in this coupling to the peak GS phase was related to memory retention. For each participant, coupling strength was quantified as the proportion of reactivation events occurring within a window centered on the peak of the group-level phase distribution (60° window, from –30° to +30° around the peak). We then computed Pearson’s correlation between the coupling strength (proportion of reactivation events around the GS peak) and memory retention (associative forgetting). In addition, to test potential effects of TMS on the relationship between reactivation and GS phase, we performed a two-sample Kuiper test to compare the GS-phase distributions between the two TMS groups and assessed the association between coupling strength and memory retention using Pearson’s correlation within each group separately.

In addition to the GS phase, we assessed time-varying changes in the GS amplitude during post-encoding rest across minute-long windows, linked with fluctuations in arousal or vigilance (Gonzalez-Castillo et al., 2022; Wong et al., 2013). To do so, we used a sliding window approach with a window size of 1 min (i.e., 30 TRs) and no overlap (Fig. 4C; Gonzalez-Castillo et al., 2022). Following prior work (Wong et al., 2013), we calculated the GS amplitude for each time window as the standard deviation of the GS percent signal change time-series (i.e., the mean-centered GS time course divided by its mean).

To determine whether reactivation occurrence co-varies with GS amplitude, we examined the relationship between reactivation event rates and GS amplitudes across the 1-min time windows using two methods: a median-split analysis and a graded correlation analysis. For the median-split approach, the 27 time windows of each participant (across three post-encoding scans) were split into two halves (i.e., 13 windows for each) according to the median GS amplitude. We then calculated the mean reactivation event rate separately within high (above the median) and low (below the median) GS-amplitude windows for each participant (Fig. 4C) and compared the two using a paired t-test. To test whether links between reactivation and GS amplitude varied with TMS, we performed a mixed ANOVA on reactivation event rates using factors of TMS site (LOC vs. Control) and GS-amplitude time window (high vs. low).

To probe more fine-grained relationships, as opposed to a median split, we measured graded relationships between reactivation and GS amplitude by computing Spearman’s correlation between the reactivation event rate and GS amplitude across time windows in each participant. Group-level significance was then tested by comparing the Fisher z-transformed correlation coefficients against 0 using a one-sample t-test. We further tested whether the degree of reactivation–GS amplitude coupling (Fisher z-transformed correlations) differed between the two TMS conditions using an independent-samples t-test.

To test whether reactivation is similarly or differentially related to memory based on the ongoing GS amplitude, which has been previously related to arousal or vigilance levels, we measured Pearson’s correlation between mean reactivation event rate and memory retention (associative forgetting) separately for high and low GS amplitude windows. We also assessed whether time windows of varying GS amplitudes are differentially related to memory using a permutation analysis. Specifically, we randomly shuffled the condition labels for the mean reactivation event rates across the high and low GS amplitude windows and recomputed, for each permutation, the difference in Pearson’s correlation coefficients with forgetting between the two GS-amplitude conditions. The observed correlation difference was then compared to the permuted null distribution (10,000 iterations). For the graded correlation approach, we used Pearson’s correlation to test whether the degree of reactivation-GS amplitude coupling (i.e., Spearman’s correlation between reactivation rate and GS amplitude) was related to associative forgetting.

To assess whether the behavioral relevance of reactivation-GS amplitude coupling differs across TMS groups, we conducted regression analyses on memory retention (associative forgetting) using regressors of TMS condition, GS amplitude measure (mean event rate in high amplitude windows for the median-split approach or the Spearman correlation for the graded correlation approach), and their interaction.

### Single Region Reactivation Analysis

Lastly, we asked whether reactivation events identified in single regions exhibit similar or distinct patterns of coupling with brain state dynamics. To address this question, we performed the brain state strength, GS phase, and GS amplitude analyses separately for reactivation events in each region, and then compared how region-specific reactivation events are differentially coupled with global brain dynamics. For the brain state strength, we ran a two-way repeated measures ANOVA with factors of brain state (four states) and reactivation region (hippocampus vs. LOC), followed by pairwise comparisons between the two regions with FDR correction. For the GS phase analysis, we used a two-sample Kuiper test to compare the phase distributions of hippocampal and LOC reactivation events, followed by separate Rayleigh tests for each region. For the GS amplitude analysis, we performed a paired t-test on the Spearman correlation (i.e., the relationship between GS amplitude and reactivation event rate across time windows within individuals) for hippocampal and LOC reactivation events, as well as separate one-sample t-tests for each region.

## Acknowledgements

This work was supported by National Institute of Health grants P50MH109429, R01MH63901, F32MH106280, R01MH139349, RF1MH128696, and R01MH138766.

## Data Availability

The data supporting the findings of this study are available from the corresponding author upon reasonable request.

## Code Availability

Custom code used for the analyses reported in this study will be made available upon publication.

## Author Contributions

AT and MD designed the original experiment. AT collected the data. AT and HWK conceptualized the present study. HWK, TX, KHN, and AT contributed to data analysis. AF, SC, and MM provided methodological and analytical inputs. HWK and AT drafted the manuscript, with feedback from all authors. All authors approved the final manuscript for submission.

## Ethics, Consent to Participate, and Consent to Publish

Ethics approval: This study was approved by the Committee for Protection of Human Subjects at the University of California, Berkeley (protocol number 2010-02-781).

Consent to participate: All participants provided informed consent prior to participation.

Consent to publish: Not applicable; the manuscript does not contain identifiable images or information from any participant.

## Supplementary Results

### TMS Modulation of Brain State-Memory Relationships during Reactivation

We further characterized whether relationships between brain state strength during reactivation and associative forgetting varied across TMS conditions. For each brain state, we asked whether associative forgetting was predicted by brain state strength during reactivation, TMS site (mean centered), and their interaction, followed by correlations within each TMS group. For State 2, consistent with the results across all participants, State 2 strength during reactivation was significantly related to forgetting when TMS site was included in the model (main effect: *b* = 1.09 ± .34, *t* = 3.20, *p* = .002). The TMS × State 2 interaction was not significant (*b* = –0.19 ± .68, *t* = –0.29, *p* = .776), indicating that the relationship between forgetting and State 2 strength during reactivation did not differ reliably across TMS sites. Indeed, relationships were in the same direction in both TMS conditions, with greater expression of the 2A pole during reactivation being associated with greater forgetting (Control TMS group, *r* = 0.51, *p* = .008; LOC TMS group, *r* = 0.35, *p* = .083; Fig. S1B).

A distinct pattern was observed for State 4. While the main effect of State 4 strength during reactivation was not significant when TMS site was modeled (*b* = –.29 ± .42, *t* = –0.70, *p* = .486), the TMS × State 4 strength interaction was significant (*b* = 2.07 ± .84, *t* = 2.48, *p* = .017), indicating that the relationships between State 4 strength during reactivation and forgetting differed between TMS sites. Specifically, this relationship emerged only in the control TMS group (*r* = –0.49, *p* = .010; Fig. S1B, top right) but was not reliable in the LOC TMS group (*r* = 0.21, *p* = .305; Fig. S1B, bottom right).

State 1 strength during reactivation was not significantly related to forgetting (main effect: *b* = 0.83 ± .47, *t* = 1.75, *p* = .086; Control TMS group: *r* = 0.19, *p* = .350; LOC TMS group: *r* = 0.29, *p* = .157; Fig. S1B, left) and this relationship did not differ between TMS sites (interaction effect: *b* = 0.66 ± .95, *t* = 0.69, *p* = .491), consistent with the prior analysis across all participants. For State 3, the main effect of state strength was not significantly related to forgetting (main effect: *b* = 0.40 ± .38, *t* = 1.06, *p* = .295), but there was a marginal TMS × State 3 interaction (*b* = –1.51 ± .76, *t* = –1.99, *p* = .052), indicating that the relationship between State 3 strength during reactivation and forgetting showed a trend-level difference across TMS sites. Greater expression of the 3A pole during reactivation was associated with increased forgetting in the Control TMS group (*r* = 0.42, *p* = .035), while this relationship was not reliable in the LOC TMS group (*r* = –0.13, *p* = .528; Fig. S1B, second right).

Together, these results indicate that the relationship between State 2 expression during reactivation and memory was relatively consistent across TMS sites, whereas the State 4 relationship differed across TMS sites and was only present in the Control TMS group; State 3 showed a similar but weaker pattern of TMS-dependent modulation. Thus, the relationship between brain state expression during reactivation and memory was selectively modulated by TMS, with one remaining stable (State 2) and other being altered by perturbation of post-encoding activity (States 3 and 4).

### Brain States and Global Signal Dynamics Associated with LOC and Hippocampal Reactivation: Full Results

To assess whether global brain dynamics linked to coordinated reactivation events (reported in the Results) reflect properties specific to coordinated hippocampal-cortical co-reactivation, or whether they generalize to reactivation events identified in individual brain regions, we examined coupling with global brain dynamics separately for hippocampal and LOC reactivation events. This approach allowed us to test the prediction that reactivation propagates from the hippocampus outward to neocortical regions along the principal gradient (Kaefer et al., 2022). This framework predicts that distinct brain state profiles would be active during hippocampal vs. LOC reactivation events, such that hippocampal reactivation is linked with brain states reflecting the association pole of the S-A axis (3B), while LOC reactivation is linked with the sensorimotor pole of the S-A axis (3A).

Indeed, reactivation events in these two regions were associated with distinct brain states (Fig. S4A). A two-way repeated-measures ANOVA on the brain state strength measure revealed significant main effects of brain state (*F*_3,153_ = 12.64, *p* < .001) and reactivation region (*F*_1,51_ = 8.87, *p* = .004), as well as a significant interaction between the two factors (*F*_3,153_ = 53.01, *p* < .001). Consistent with the principal gradient prediction, State 3 strength significantly differed between hippocampal versus LOC reactivation (pairwise comparison: *t_51_*= –9.05, *p_FDR_* < .001). LOC reactivation events were coupled with the 3A brain state pole (one sample t test: *t_51_* = 9.98, *p_FDR_* < .001), indicating LOC reactivation is more closely aligned with the sensorimotor pole of the S-A axis (Margulies et al., 2016). By contrast, hippocampal reactivation tended to occur during the 3B brain state pole (*t_51_* = –2.32, *p_FDR_* = .024), reflecting the association pole of the S-A axis. In addition to State 3, State 1 strength was greater or biased towards the A pole during hippocampal compared to LOC reactivation (pairwise comparison: *t_51_* = 7.32, *p_FDR_* < .001). This indicates that hippocampal reactivation was more reliably coupled with an internally oriented brain activity pattern (1A, one sample t test: *t_51_* = 9.96, *p_FDR_* < .001), strongly weighting the DMN. In contrast, LOC reactivation was not reliably coupled with State 1 (*t_51_* = –1.21, *p_FDR_*= .231), suggesting that visual reactivation in isolation does not preferentially occur during internally or externally oriented states, instead being linked with activation of the broader visual network (evident in States 2 and State 3). In contrast to States 1 and 3, which were distinctly linked with hippocampal vs. LOC reactivation events, States 2 and 4 did not reliably differ in their coupling to hippocampal vs. LOC reactivation (pairwise comparisons: State 2: *t_51_* = –0.55, *p_FDR_* = .587; State 4: *t_51_* = 0.06, *p_FDR_* = .952), suggesting that coupling of coordinated hippocampal-LOC reactivation with States 2 and 4 reflects a shared property, rather than being driven by a specific region. Together, these findings align with predictions of the gradient cascade model of reactivation (Kaefer et al., 2022) and additionally reveal that hippocampal reactivation is more selectively tied to an internally oriented processing state, suggesting that the conditions supporting hippocampal reactivation may be more constrained than those supporting sensory cortical reactivation.

Analysis of the GS showed that reactivation events in the two regions differed in their preferred GS phase (Kuiper two-sample test: *V* = 0.072, *p* < .001; Fig. S4B). While hippocampal reactivation events were associated with the GS peak (Rayleigh test: *p* < .001), consistent with coordinated events (Fig. 4B), LOC events were linked with earlier GS phases, peaking around –45° (Rayleigh test: *p* < .001; Fig. S4B). This observation of a phase offset, with LOC reactivation preceding hippocampal reactivation in the GS phase, appears inconsistent with the gradient cascade model, which predicts that hippocampal reactivation would precede cortical reactivation.

Lastly, we asked whether LOC and hippocampal reactivation events occurred more frequently during minute-long time windows with high GS amplitude. Indeed, analogous to coordinated reactivation events, reactivation rate in each region was positively correlated with GS amplitude across time windows (i.e., Spearman correlation greater than 0; one-sample t-tests: LOC: *t_51_* = 3.92, *p* < .001; hippocampus: *t_51_* = 8.84, *p* < .001). However, this link between reactivation and GS amplitude was significantly stronger for hippocampal reactivation (paired t-test: *t_51_* = 3.54, *p* < .001; Fig. S4C). This observation of stronger coupling between hippocampal reactivation and GS amplitude, combined with differential locking of hippocampal reactivation to the peak GS phase (reported above), suggests a tighter association between hippocampal reactivation and GS fluctuations linked to lower arousal.

---

**Figure S1.**
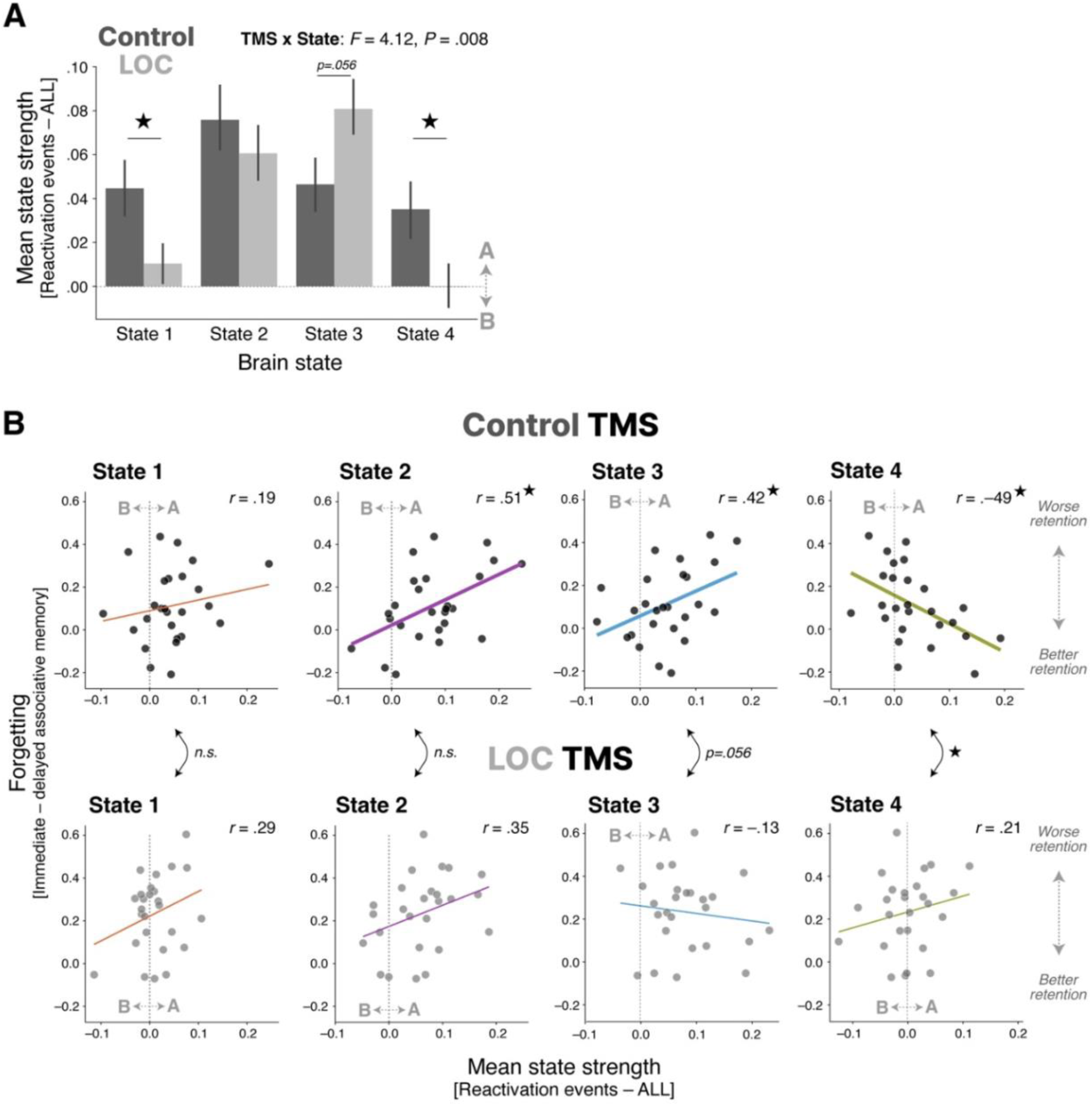
TMS effects on brain state coupling with reactivation and its link to memory. (A) Mean brain state strength during reactivation events, relative to mean strength across all time points, shown separately for Control TMS (dark gray) and LOC TMS (light gray) groups. TMS to LOC reduced State 1 and 4 strengths (expression of the 1A and 4A poles) during reactivation relative to Control TMS, and marginally increased State 3 strength (3A expression) during reactivation. Stars indicate significant pairwise comparisons (p < .05). (B) Scatter plots show the relationship between brain state strength during reactivation and forgetting (immediate minus delayed associative memory score) for each of the four brain states, plotted separately for the Control TMS (top) and LOC TMS (bottom) groups. Each data point denotes a single participant. Stars indicate significant correlations or interactions by TMS (p < .05).

**Figure S2.**
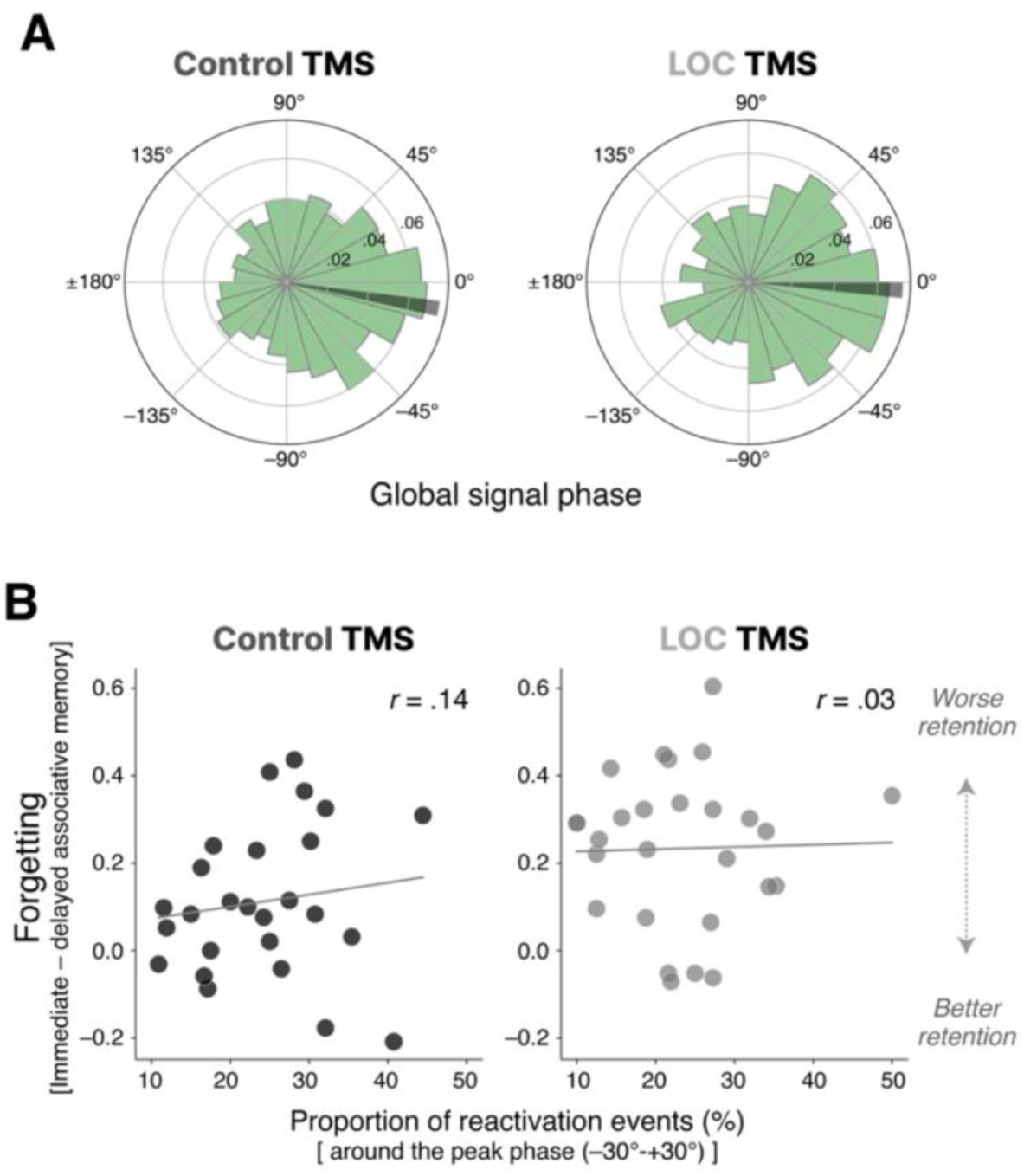
TMS effects on global signal (GS) phase coupling with reactivation and its link to memory. (A) Distribution of GS phase of reactivation time points, aggregated across participants, shown separately for the Control TMS (left) and LOC TMS (right) groups. Reactivation events clustered around the peak GS phase (0°) in both groups, and this phase-locking did not differ by TMS site. (B) Scatter plots show the relationship between the proportion of reactivation events occurring near the peak GS phase (−30° to +30°) and forgetting (immediate - delayed associative memory score), plotted separately for the Control TMS (left) and LOC TMS (right) groups. Each data point denotes a single participant.

**Figure S3.**
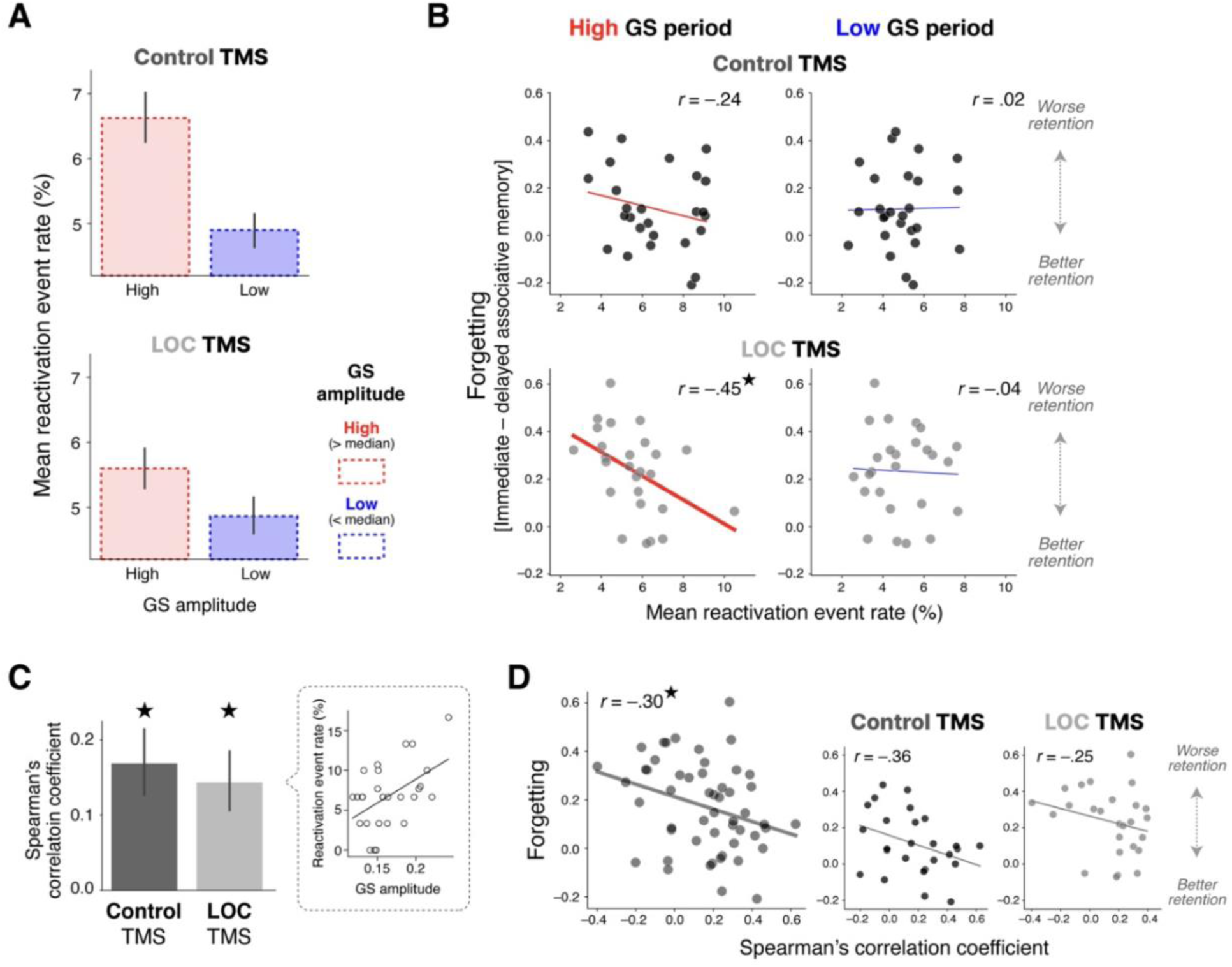
TMS effects on global signal (GS) amplitude coupling with reactivation and its link to memory. (A) Mean reactivation event rate in high-versus low-amplitude GS windows, shown separately for the Control TMS (top) and LOC TMS (bottom) groups. (B) Scatter plots show the relationship between mean reactivation event rate and forgetting (immediate - delayed associative memory score) for high-GS (left) and low-GS (right) amplitude periods, plotted separately for the Control TMS (top) and LOC TMS (bottom) groups. Each data point denotes a single participant. (C) Spearman’s correlation between GS amplitude and reactivation event rate across time windows within individual participants, shown separately for the Control TMS and LOC TMS groups (stars indicate correlations significantly above 0, p < .05). Inset shows an example scatter plot of reactivation event rate against GS amplitude across time windows for one participant. (D) Scatter plots show the relationship between the Spearman’s correlation coefficient (GS amplitude reactivation rate) and forgetting, across all participants (left) and separately for the Control TMS (middle) and LOC TMS (right) groups. Each data point denotes a single participant.

**Figure S4.**
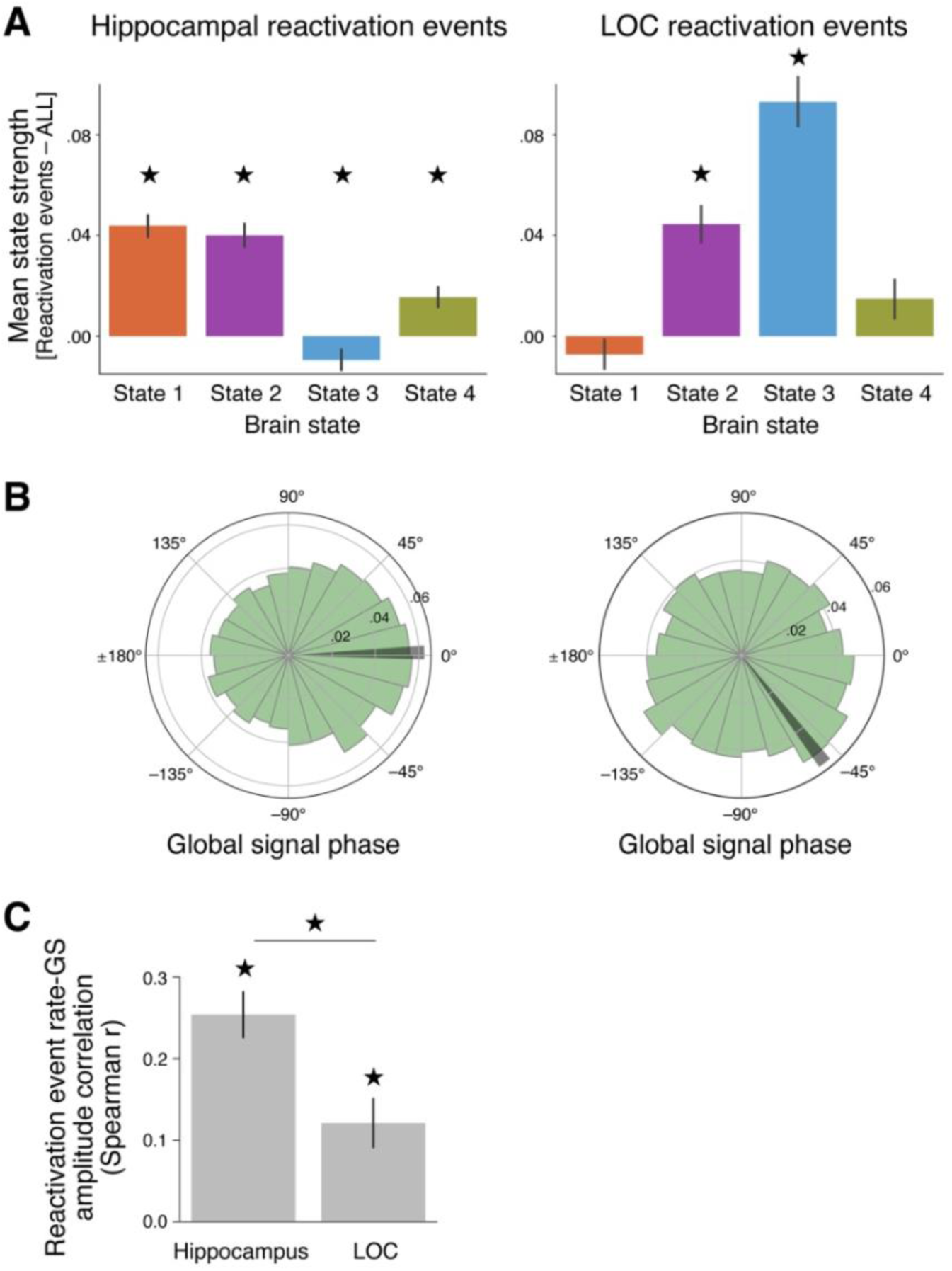
Region-specific coupling of hippocampal and LOC reactivation with brain states and global signal (GS) phase. **(A)** Mean brain state strength during reactivation events, relative to mean strength across all time points, for hippocampal (left) and LOC (right) reactivation events. Hippocampal reactivation was coupled with States 1, 2, and 4, and with the ‘B’ pole of State 3, whereas LOC reactivation was coupled with States 2 and 3 (biased toward the ‘A’ pole) but not reliably with State 1. Stars indicate FDR-corrected one-sample t-tests against 0 (p < .05). (B) Distribution of GS phase of reactivation time points for hippocampal (left) and LOC (right) reactivation events. Hippocampal reactivation clustered around the peak GS phase (0°), whereas LOC reactivation was linked to an earlier GS phase (peaking around ∼45°), indicating a phase offset between the two regions. (C) Spearman’s correlation between reactivation event rate and GS amplitude across time windows within participants, shown separately for Hippocampal and LOC reactivation events (stars indicate correlations significantly above 0, and significant differences in correlation between the two regions, p < .05).

